# Pitavastatin counteracts venetoclax resistance mechanisms in acute myeloid leukemia by depleting geranylgeranyl pyrophosphate

**DOI:** 10.1101/2025.10.27.684888

**Authors:** Roberta Buono, Dennis Juarez, Madhuri Paul, Sarah J. Skuli, Ian B. Wong, Ishita Tarnekar, Zhili Ying, Izabelle Le, Gerald Wertheim, A’ishah Bakayoko, Marisa Kruidenier, Said M. Sebti, Marina Konopleva, Angela G. Fleischman, Cholsoon Jang, Martin Carroll, David A. Fruman

**Author notes:** Address Correspondence: David A. Fruman, PhD, UC Irvine, 839 Health Sciences Road, 216 Sprague Hall, Irvine, CA, 92697-3905. or Roberta Buono, PhD, UC Irvine, 839 Health Sciences Road, 216 Sprague Hall, Irvine, CA, 92697-3905. these authors contributed equally to this work (Roberta Buono, Dennis Juarez, Madhuri Paul). Competing Interests Statement: SMS is co-inventor of GGTI-2418 and GGTI-2417, and is Scientific Founder of Prescient Therapeutics. MK receives research support and serves a consultant and advisory board member for AbbVie. The other authors have no competing interests to disclose.

## Abstract

The BCL2 inhibitor venetoclax has therapeutic activity in several hematological malignancies. In acute myeloid leukemia (AML), venetoclax combined with hypomethylating agents is the standard of care for patients unfit for intensive chemotherapy, but intrinsic and acquired resistance are common. Loss of p53 function is strongly associated with venetoclax resistance, and adding venetoclax to 5-azacitidine provides no overall survival benefit in *TP53*-mutant AML. Other frequent mechanisms of venetoclax resistance in AML include *FLT3* mutations, MCL-1 upregulation, and altered mitochondrial metabolism. Unfortunately, it has been challenging to develop agents that target these mechanisms directly and combinatorially. Here we report that pitavastatin, an inhibitor of HMG-CoA-reductase, promotes apoptosis and overcomes several venetoclax resistance mechanisms in human AML cells. At clinically achievable concentrations, pitavastatin treatment has potent cytotoxic activity in cells with mutations in *TP53* or *FLT3*. The apoptotic mechanism involves p53-independent PUMA upregulation and reduced MCL-1 expression. Pitavastatin also suppresses mitochondrial gene expression and oxidative metabolism. The pro-apoptotic actions of pitavastatin depend on depletion of geranylgeranyl pyrophosphate (GGPP) and can be recapitulated by inhibiting GGPP synthase or geranylgeranyltransferase-1 enzymes. These results provide a mechanistic rationale for adding pitavastatin to AML regimens to prevent or overcome venetoclax resistance.

## Introduction

B cell lymphoma 2 (BCL2) is an oncoprotein and a member of a family of apoptosis-regulating proteins (1). The pro-survival family members (BCL2, BCL-xL, MCL-1, BCL-w, BFL-1) contain four domains with homology to BCL2 (BH1, BH2, BH3, BH4). Levels of these proteins are often elevated in blood cancer cells to escape stress-induced increases of pro-apoptotic proteins, including “BH3-only” proteins such as BIM and PUMA (1–3). BH3 mimetics are cell-permeable small molecules that block the BH3-binding pocket of BCL2 family members, inhibiting their pro-survival function and promoting apoptosis (1–3). Their clinical utility is validated by BCL2-selective venetoclax (ABT-199) regimens for front-line treatment of chronic lymphocytic leukemia (CLL) and acute myeloid leukemia (AML) (4,5). However, there remains a critical need to identify combinations that further broaden and deepen responses to venetoclax and prevent the emergence of resistance (6).

AML demonstrates diverse resistance mechanisms to BCL2 inhibition. Despite initial success in the treatment of *TP53*-mutant CLL (4), subsequent basic and clinical studies highlight that bi-allelic *TP53* loss is a frequent mechanism of resistance in AML (7–9). Indeed, while venetoclax combined with 5-azacitidine (AZA) extends survival in AML patients unfit for intensive chemotherapy (10), this combination provides no overall survival benefit compared to AZA alone in patients with *TP53* loss (11,12). Other factors associated with venetoclax resistance in AML include reduced mitochondrial priming through upregulation of other anti-apoptotic proteins such as MCL-1 (13), certain genetic drivers (e.g., *FLT3*-internal tandem duplication (ITD) and *RAS*; (8,14)), *BCL2* mutations (15), and altered mitochondrial dynamics including increased respiration (16–19).

The mevalonate pathway is a targetable cancer dependency that supports proliferation and survival (20,21). Many oncogenes increase cellular demand for mevalonate pathway outputs, while driving mevalonate flux by increasing expression of HMG-CoA-reductase (HMGCR) or other pathway enzymes (21). This has spurred interest in repurposing statins (selective HMGCR inhibitors) for oncology (21–24). We reported that statins promote apoptosis and sensitize to venetoclax in blood cancer cell lines and primary cells (25,26), a finding independently validated by others (27,28). Our retrospective analysis of venetoclax clinical trials revealed that statin use is associated with significantly increased rates of complete response in CLL or multiple myeloma (MM) (25,26). These findings provided rationale for a prospective phase 1 clinical trial (clinicaltrials.gov identifier NCT04512105) testing pitavastatin in combination with venetoclax in AML and CLL. This trial found that the addition of pitavastatin to venetoclax regimens is tolerable (29).

Barriers that have limited the clinical translation of statins as oncology drugs include the many mechanisms of action described in the literature and the underappreciated role of statin pharmacology in design of clinical trials (21,30). We previously determined that the pro-apoptotic effect of statins is due to depletion of cellular pools of geranylgeranyl pyrophosphate (GGPP) (25,26), the isoprenoid substrate for protein geranylgeranylation, and this finding is generalizable to several of the described mechanisms of statin’s anti-cancer effects (21,31).

Concerning statin pharmacology, we found that pitavastatin has approximately 3-fold greater potency than other statins in cell-based assays, and acts at clinically achievable concentrations to block geranylgeranylation and promote apoptosis in MM cells (26). Pitavastatin in our phase I trial also demonstrated preliminary signs of efficacy with all subjects achieving complete remission (29).

Here we assess statin efficacy against AML venetoclax resistance using cell line models and primary cells. To focus on mevalonate pathway outputs relevant to survival signaling in AML cells, we identify cellular responses that are induced by pitavastatin in a manner dependent on GGPP depletion and are recapitulated by geranylgeranyltransferase-1 (GGTase-1) inhibitors.

These analyses revealed that pitavastatin sensitizes to venetoclax via p53-independent upregulation of PUMA and by rewiring cellular transcription to disrupt mitochondrial gene expression and metabolism. In a subset of AML cells, pitavastatin diminishes the expression of MCL-1. Together, these results show that pitavastatin can overcome several common mechanisms of venetoclax resistance in AML and should be assessed in combination with venetoclax and AZA in future clinical trials.

## Materials and Methods

### Additional Methods provided in the supplement

#### Cell Culture

AML cell lines (Table 1) were obtained from commercial repositories (ATCC or DSMZ) and authenticated annually through STR profiling using the University of Arizona Genomics Core. MOLM13 parental and venetoclax resistant were described previously (32). Cell lines were tested for mycoplasma infection annually. Unless otherwise noted, cell lines used in the study were maintained in RPMI-1640, 10% FBS, 1% penicillin-streptomycin-glutamine (Gibco) and 10 mM HEPES (Corning).

Human cells were provided by UCI biobank or U-Penn (Table 2) and cultured in IMDM supplemented with 15% BIT 9500 with and 1 µM StemRegenin 1 (Stemcells Technologies), plus human cytokines (Peprotech): SCF (100 ng/ml), Flt3-Ligand (50 ng/ml), IL-3 (20 ng/ml), G-CSF (20 ng/ml), 1:100 penicillin-streptomycin-glutamine, and 50 µM beta mercaptoethanol.

**Table 1.**
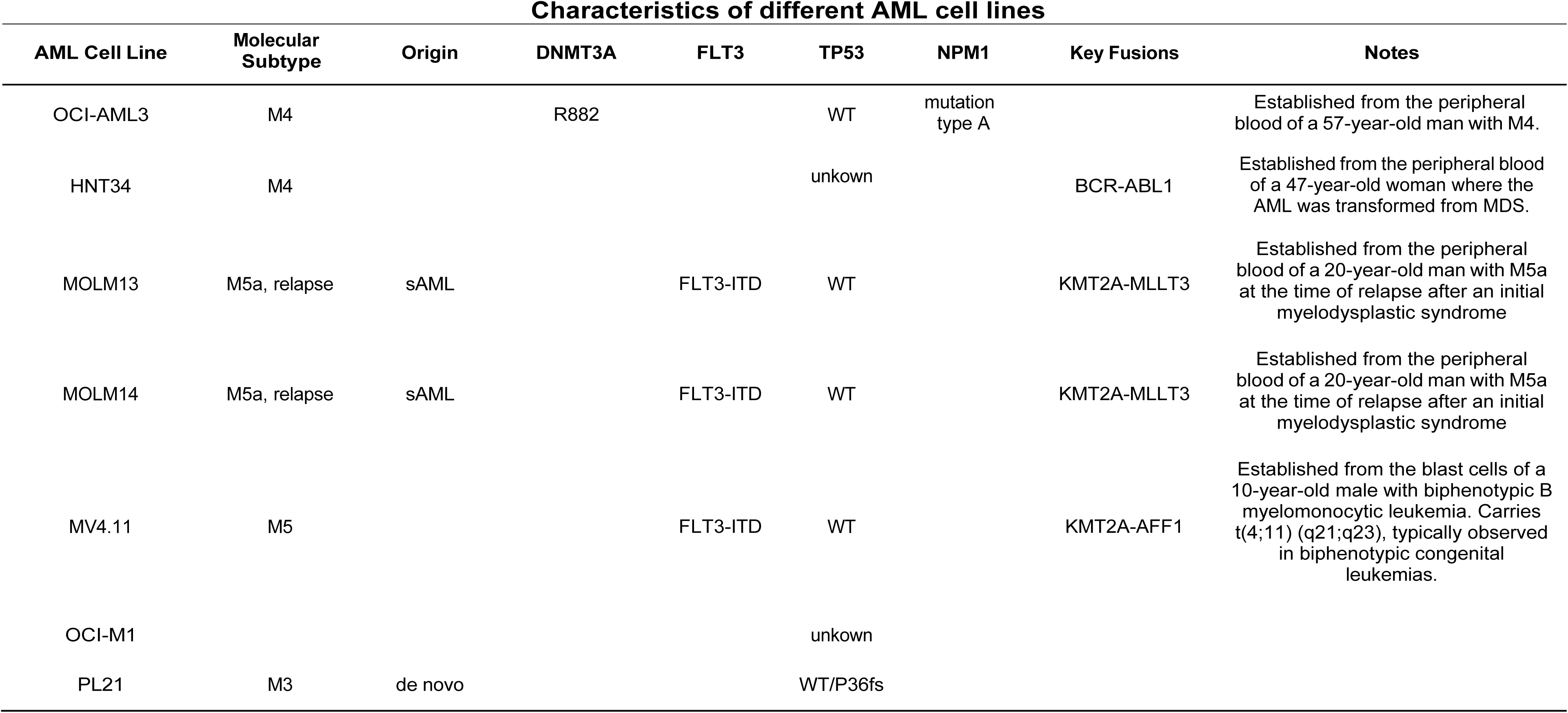

#### Statistics and experimental design

Statistical analysis was performed in Prism10 using one-way or two-way ANOVA, as appropriate for each experiment and indicated in the legends. Post-tests for multiple comparisons were recommended by Prism and applied as noted in the Figure legends. All samples represent biological replicates. Unless otherwise specified in figure legends, all center values shown in graphs refer to the mean. The number of biological replicates (n) and details of the tests are provided in the figure legends. p < 0.05 was considered statistically significant and was annotated throughout as * p < 0.05, ** p < 0.01, *** p < 0.001 and **** p < 0.001.

## Results

### Pitavastatin induces p53-independent cytotoxicity at sub-micromolar concentrations in AML cells

Previously we reported that simvastatin sensitizes AML cell lines to venetoclax cytotoxicity (25), and that pitavastatin is cytotoxic in MM cell lines at lower concentrations than simvastatin (26). Here we tested pitavastatin in a panel of AML cell lines with different genetic lesions (Table 1). Cytotoxicity assays demonstrated that 48-hour treatment with pitavastatin concentrations as low as 300 nM reduced viability in OCI-AML3 and HNT-34 cells and enhanced the weak cytotoxic effect of venetoclax (Figure 1A). A similar effect was observed in MV4-11 cells at 1 µM pitavastatin (Figure 1A). The mechanism by which pitavastatin enhanced cytotoxicity was caspase-dependent, as it was blocked by the inhibitor Q-VD-OPh at 24hr (Fig. S1A). In the MOLM13 and PL-21 cell lines, pitavastatin alone had minimal cytotoxic activity but enhanced the effect of venetoclax (Figure 1A). Pitavastatin retained cytotoxicity in a venetoclax-resistant subline of MOLM13 (32) (Fig. S1B). The combination of pitavastatin and venetoclax was synergistic in OCI-AML3, HNT-34, MV4-11, and MOLM13 cells (Table S1; the latter three cell lines have *FLT*3-ITD). We identified a pitavastatin-resistant AML cell line (OCI-M1) (Figure 1A, Table S1) that we used as a comparator for mechanistic studies. Consistent with prior publications (25), simvastatin enhanced venetoclax cytotoxicity but only at concentrations above 1 µM in OCI-AML3 and MOLM13 cells (Fig. S1C).

**Figure 1:**
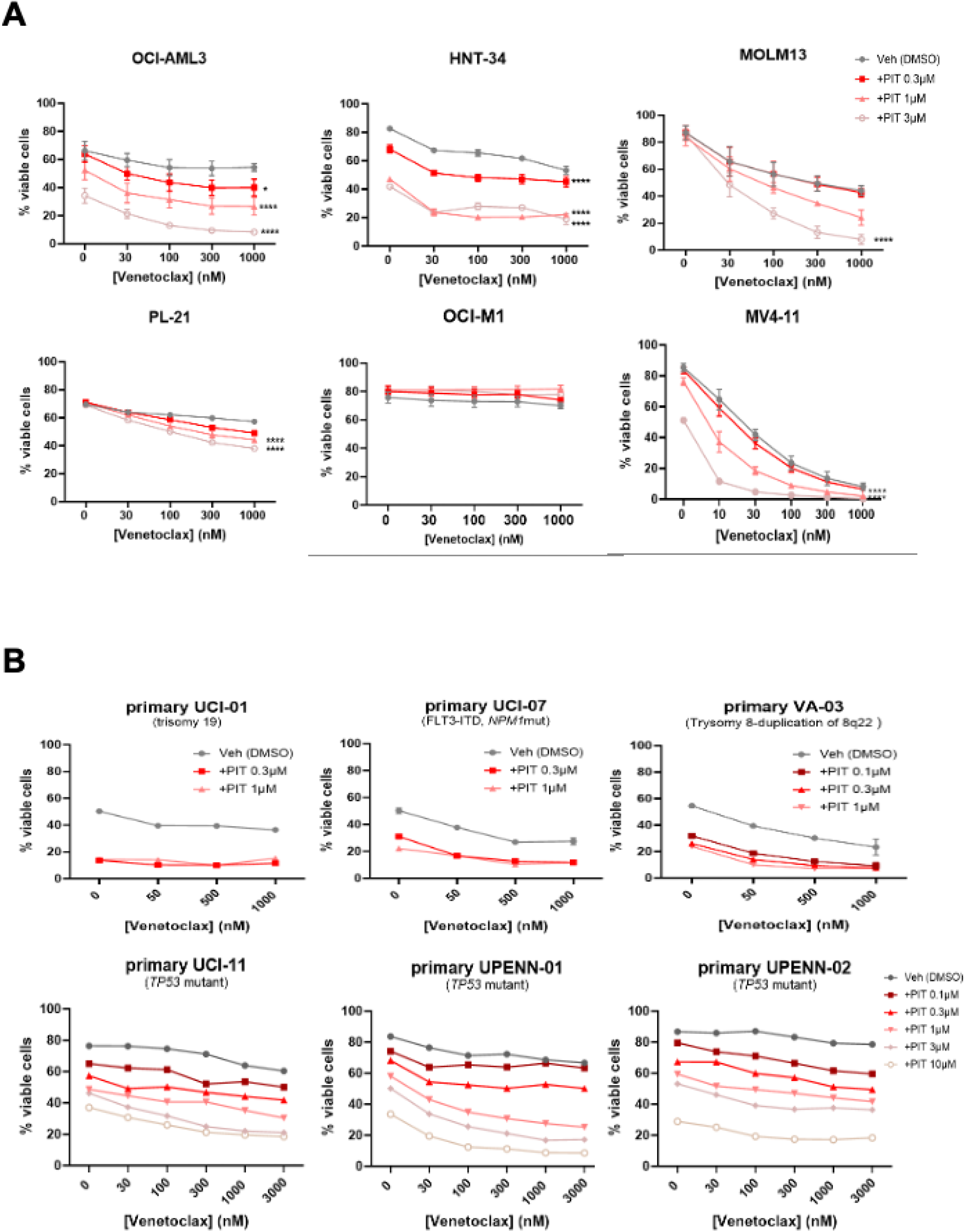
Pitavastatin has potent cytotoxicity in venetoclax-resistant AML cell lines and in patient-derived AML cells. (A) AML cell lines were treated for 48hr with various concentrations of venetoclax alone or with pitavastatin (PIT) at 0.3, 1 and 3 µM. Data are expressed as mean +/-SEM *p < 0.05, ****p < 0.0001. Two-way ANOVA with Tukey’s post hoc test. (B) Patient-derived AML myeloblasts cultured in media with supportive cytokines were treated for 48hr with titrated concentrations of PIT and venetoclax as indicated. For both panels the percentage of viable cells was determined by flow cytometry using Annexin V and PI staining. Identified genetic lesions are shown in parentheses – note three *TP53* mutant samples on the bottom row were all pitavastatin-sensitive.

We extended these studies to cultures of primary AML cells obtained from cryopreserved adult AML mononuclear specimens (Table 2). Primary cultures were more pitavastatin sensitive than established cell lines with notable cytotoxicity at 100 – 300 nM including three samples with *TP53* mutations and three out of five samples with *FLT3*-ITD (Figure 1B and S2A, B, C).

Venetoclax further enhanced the killing of several of the primary AML samples (Figure 1B). In addition, pitavastatin increased cytotoxicity of the combination of venetoclax and AZA in several primary AML samples (Fig. S2C). In most of these samples, the combination of venetoclax plus pitavastatin was more effective than venetoclax plus AZA.

We showed previously that expression of a dominant-negative p53 did not affect simvastatin-mediated cytotoxicity or venetoclax sensitization in OCI-AML3 cells (25). To investigate further the relationship of p53 status to pitavastatin sensitivity, we used isogenic MOLM14 cell lines with either wild-type or bi-allelic deletion of *TP53*, generated by CRISPR/Cas9 (33). In parental MOLM14 cells, pitavastatin in the 1-3 µM concentration range was sufficient to promote cell death (Fig. S3). The p53-deficient MOLM14 variants, which are resistant to cytarabine (33) and less sensitive than control MOLM14 cells to venetoclax (Fig. S3), remained sensitive to pitavastatin and to the synergistic or additive cytotoxicity of pitavastatin plus venetoclax (Fig. S3, Table S2).

Overall, the findings that pitavastatin at submicromolar concentrations (clinically achievable with FDA-approved doses, enhances the cytotoxic activity of venetoclax in a variety of AML cell lines and primary cells, and that primary AML cells are more sensitive to pitavastatin at lower concentrations than AML cell lines, suggest that addition of pitavastatin to venetoclax regimens in AML may be a feasible approach for treatment, including in cases with p53 inactivation or *FLT3*-ITD.

### Pitavastatin alters expression of BCL2 family members PUMA and MCL-1

p53-independent upregulation of PUMA is a general mechanism by which statins sensitize blood cancer cells to BH3 mimetics (25,26). Consistent with our previous findings, pitavastatin treatment increased expression of PUMA protein in OCI-AML3, HNT-34 and MV4-11 cell lines that are sensitive to statin cytotoxicity (Fig. S4A, B, C). Pitavastatin treatment caused lesser or no PUMA upregulation in cell lines that are less sensitive (MOLM-13, PL-21) or resistant to cytotoxicity (OCI-M1) (Fig. S4A, B, D). Additionally, in isogenic *TP53* deleted MOLM14 cells, pitavastatin induced PUMA expression to a comparable degree in control (Fig. S5A).

CRISPR/Cas9-mediated knockout of the *BBC3* gene encoding PUMA (Fig. S5B, C) partially reversed the cytotoxic effect of pitavastatin in OCI-AML3 and MOLM-14 cells but not in MOLM-13 cells (Figure 2A).

**Figure 2:**
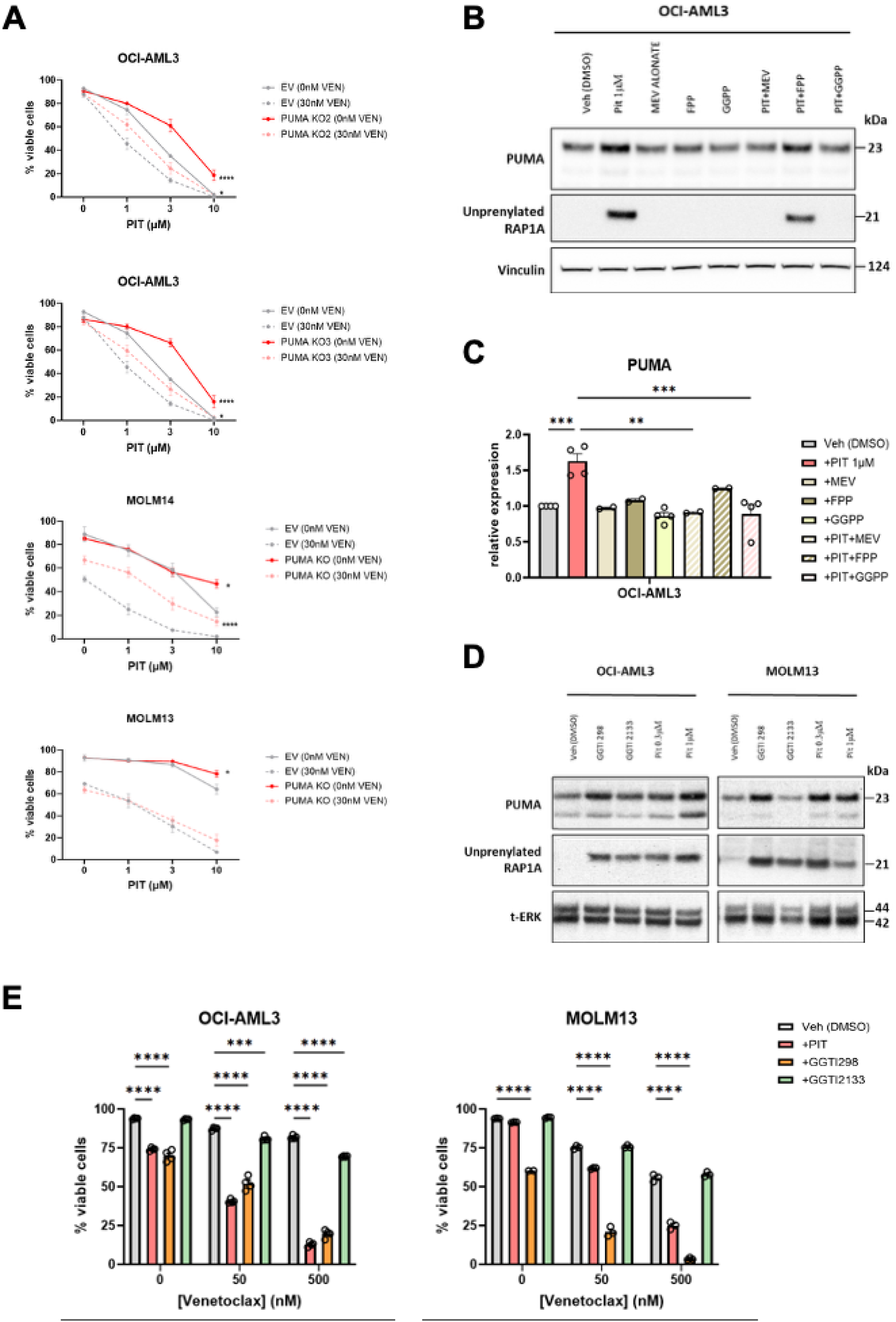
PUMA upregulation contributes to pitavastatin cytotoxicity, is dependent on GGPP depletion, and is recapitulated by inhibition of GGTase-1. (A) The cytotoxic effect of pitavastatin +/-30nM venetoclax was measured in two clones of OCI-AML3 cells with PUMA knockout and one clone each of MOLM14 and MOLM13. Two-way ANOVA with Tukey’s post hoc test, n = 3. (B) PUMA expression in a statin-sensitive AML cell line (OCI-AML3) was assessed by western blot. An antibody to vinculin was used as a loading control. Detection of unprenylated RAP1A was used to determine which treatments suppress protein geranylgeranylation. Note rescue of PIT effect on PUMA by mevalonate (MEV) and GGPP more than by FPP. (C) Quantitation of four replicates of the experiment in panel B, with PUMA expression in vehicle (Veh) treated cells set at 1. Data are expressed as mean +/-SEM ** p < 0.01, ***p < 0.001, one-way ANOVA with Tukey’s post hoc test, n = 2-4. (D-E) Different GGTase-1 inhibitors have distinct effects on PUMA expression and viability. (D) Representative western blot of PUMA expression in OCI-AML3 and MOLM13 treated with pitavastatin (0.3 or 1 µM), GGTI298 (10 µM) or GGTI2133 (10 µM). (E) Graphs showing percentage of viable OCI-AML3 and MOLM13 measured by Annexin V and PI staining after 48h of treatment with pitavastatin (1 µM+, GGTI298 (10 µM) or GGTI2133 (10 µM) in combination with venetoclax. Data are expressed as mean +/-SEM ***p < 0.001, ****p < 0.0001. Two-way ANOVA with Tukey’s post hoc test, n ≥ 3.

Increased expression of and dependence on MCL-1 can confer venetoclax resistance in AML cells (34,35). We assessed how venetoclax and pitavastatin co-treatment affects protein expression of PUMA and MCL1. In OCI-AML3, MOLM13, and MOLM14 cells, pitavastatin reduced expression of MCL-1 and/or prevented the increase in MCL-1 following venetoclax treatment (Fig. S6A, B). In general, the combination of pitavastatin with venetoclax reduced MCL-1 expression more than pitavastatin alone. We also noted that in some AML cell lines, venetoclax reduced PUMA expression and suppressed PUMA upregulation by pitavastatin (Fig. S6A, B). In the venetoclax-resistant MOLM13 subline, pitavastatin increased low basal PUMA expression but did not reduce the high basal MCL-1 expression (Fig. S6C). In two of the three *TP53* mutant primary AML cells we tested, 24hr treatment with pitavastatin reduced MCL-1 or increased PUMA expression (Fig. S6D). Thus, pitavastatin can alter the balance of pro-and anti-apoptotic BCL2 family members via at least two mechanisms.

### Pitavastatin increases cytotoxicity and PUMA expression via depletion of GGPP and inhibition of geranylgeranylation

In most blood cancer cell models, statin cytotoxicity is linked to depletion of isoprenoid intermediates downstream of mevalonate (36,37). Using an antibody that detects the unprenylated form of RAP1A GTPase, we found that pitavastatin inhibited geranylgeranylation at 1 µM or lower concentrations in all cell lines except OCI-M1 (Figure 2B, D and S4C, D). In addition, supplementing culture medium with excess GGPP prevented pitavastatin-induced PUMA expression (Figure 2B, C) and cytotoxicity (Fig. S7A) in statin-sensitive cells. This result suggests that GGPP depletion is required for the pitavastatin mechanism of action. To determine whether GGPP-depletion is sufficient, we used an inhibitor of GGPP synthase (THZ145; (38)) and found that it had cytotoxic activity in OCI-AML3 cells and increased PUMA expression to a similar extent as pitavastatin (Fig. S7B, C).

We showed previously that a GGTase-I inhibitor, GGTI-298, increased PUMA in AML cells (25). Unexpectedly, while GGTI-298 upregulated PUMA, had cytotoxic activity, and sensitized to venetoclax, the structurally related GGTI-2133 had minimal effects on these functional readouts (Figures 2D, E and S7B, C). GGTI-2133 had equivalent or greater ability to reduce prenylation of RAP1A, a commonly used readout of GGTase-1 activity (Figure 2D and S7B). Furthermore, both compounds reduced actin polymerization, which is driven by GTPases such as RHO, RAC and CDC42 known to be substrates of GGTase-1 (21,31) (Fig. S8A). Thus, GGTI-2133 and GGTI-298 both have observed activity on inhibition of GGTase-I substrates, yet GGTI-298 more closely phenocopies the anti-cancer effects of pitavastatin on AML cell function. It is unlikely that the effects of GGTI-298 are simply due to off-target activity on farnesyltransferases or other enzymes, given that GGPP synthase inhibition had cytotoxic activity similar to statins. Moreover, a more selective and potent GGTase-1 inhibitor (GGTI-2417; (39)) exerted cytotoxic activity in AML cell lines and increased PUMA expression and unprenylated RAP1A (Fig. S8B-D). Together these data support the conclusion that GGTase-1 is a key prenyltransferase mediating pro-survival activities of GGPP in AML cell lines.

### Pitavastatin and GGTI-298 elicit widespread transcriptome changes including enhanced apoptotic gene signatures

A challenge to applying transcriptome studies to decipher statin cytotoxic mechanism is that there are multiple outputs of the mevalonate pathway, of which only some play key roles in cell survival regulation. We reasoned that differentially expressed genes (DEGs) shared in pitavastatin and GGTI-298 treatment conditions but not after GGTI-2133 treatment are most relevant to the cytotoxic mechanism. In accord, we treated OCI-AML3 and HNT-34 cells with vehicle alone (0.1% DMSO) or pitavastatin (PIT; 1 µM) or GGTIs (10 µM) and harvested RNA for bulk RNA sequencing. We chose a time point (16hr) before the onset of apoptosis but when PUMA upregulation is observed at the protein level.

Pitavastatin and GGTI-298 each caused broad changes in gene expression (vs. vehicle condition, Supplementary Excel File 1) with a high degree of overlap in both OCI-AML3 (Figure 3A) and HNT-34 (Fig. S9A). GGTI-2133 treatment altered expression of many fewer genes in both cell lines (Figure 3A and S9A, Supplementary Excel File 1). Principal component analysis (PCA) supported the similarity of cells treated with GGTI-2133 or vehicle alone (Fig. S9B). In accord, unbiased transcriptome-wide analysis of rlog-based sample distances demonstrated clustering of vehicle with GGTI-2133 and of PIT with GGTI-298 in both OCI-AML3 and HNT-34 (Fig. S9C). Gene set enrichment analysis (GSEA) for Hallmark and Reactome pathways in OCI-AML3 cells showed several shared responses to pitavastatin and GGTI-298, including reduced expression of MYC and E2F targets and an increase in genes associated with apoptosis (Figure 3B). Similar gene sets were identified in HNT-34 cells, which also displayed a signature of cholesterol homeostasis and SREBP targets suggesting a stronger feedback response (Figure 3B). The *SREBF2* mRNA was increased by both pitavastatin and GGTI-298, but not GGTI-2133, with a greater effect in HNT-34 cells (Fig. S10). In agreement with the increase PUMA protein expression, the *BBC3* gene encoding PUMA was significantly upregulated by pitavastatin and GGTI-298, but not GGTI-2133, in both cell lines (Fig. S10).

**Figure 3:**
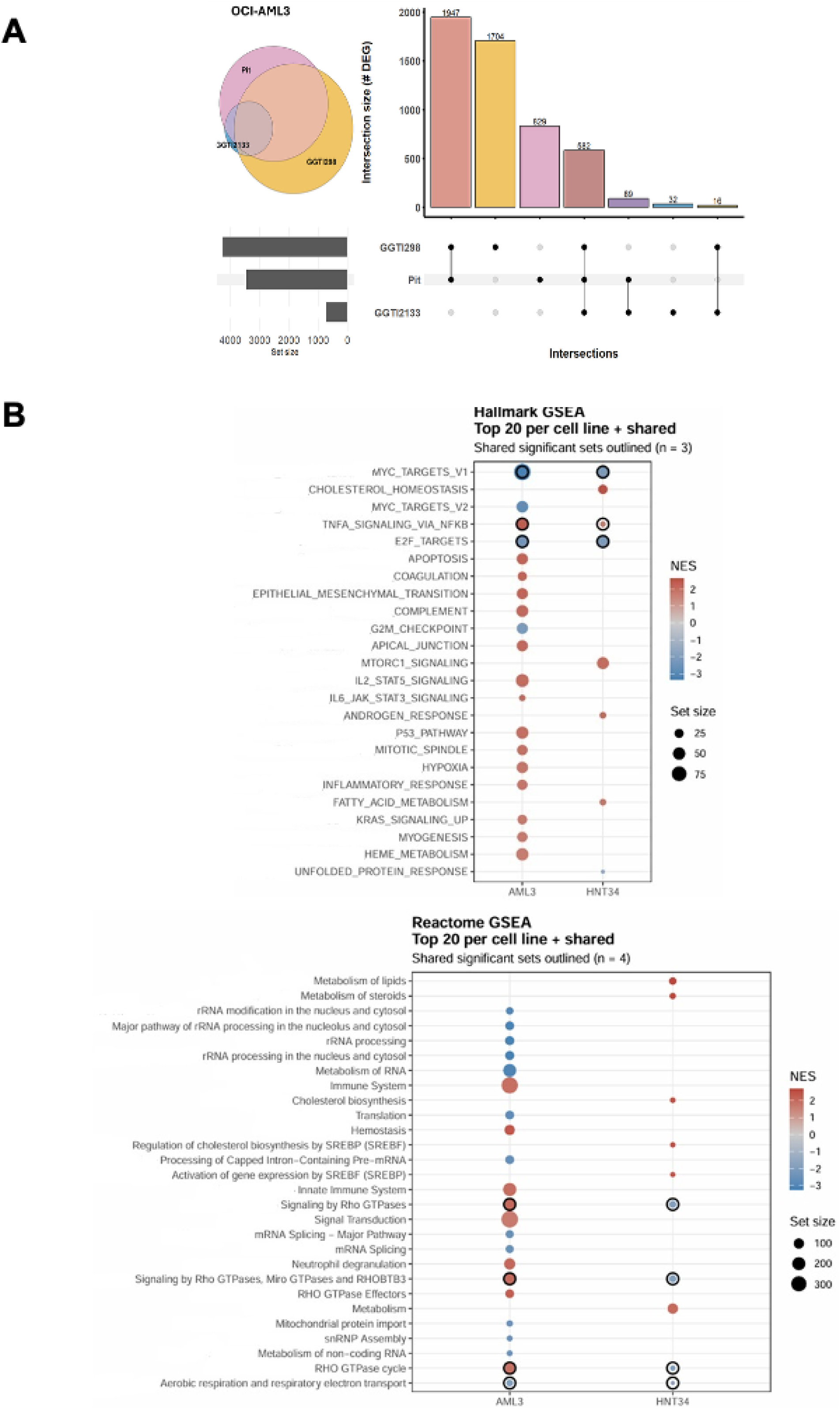
Transcriptome analysis identifies shared features of the response to PIT and GGTI-298. (A) DEGs in OCI-AML3 cells treated with PIT, GGTI-298, or GGTI-2133, relative to vehicle, were identified by DESeq2, and DEG overlaps were depicted by Venn diagrams and summarized with UpSet plots. (B) GSEA for hallmark and reactome pathways shared by treatment with PIT or GGTI-298. Red indicates GSEA pathways that are upregulated, blue downregulated. Outlined dots are pathways shared in both OCI-AML3 and HNT-34, including downregulation of MYC and E2F targets, and aerobic respiration. The diameter of each dot corresponds to gene set size.

### Transcriptome analysis of pitavastatin effects dependent on GGPP depletion

As a second approach to defining statin cytotoxic mechanism, we determined which gene expression changes induced by pitavastatin are dependent on GGPP depletion. We conducted RNA sequencing on OCI-AML3 cells treated for 16hr with vehicle alone, pitavastatin, GGPP or PIT + GGPP. Compared to vehicle-treated samples, there were ∼1400 DEGs in the PIT-treated condition but only 129 in the GGPP condition and 103 in the PIT+GGPP condition (Figure 4A). Applying a log_2_ fold-change (LFC) cut-off of +/-0.1, there were only three DEGs in the PIT+GGPP vs. Vehicle condition and one in the GGPP vs. Vehicle (Supplementary Excel File 2). PCA showed that PIT-treated samples clustered together and distant from the other three groups (Fig. S11A). These data indicate that statin-mediated transcriptional reprogramming at 16hr is almost entirely dependent on GGPP depletion. In MOLM13 cells, which are less pitavastatin sensitive than OCI-AML3 cells (Figure 1), pitavastatin caused many fewer DEGs yet these were still largely dependent on GGPP depletion (add S11B). Hallmark GSEA analysis of OCI-AML3 cells showed downregulation of MYC and E2F targets (Figure 4B), consistent with the first transcriptome experiment.

**Figure 4:**
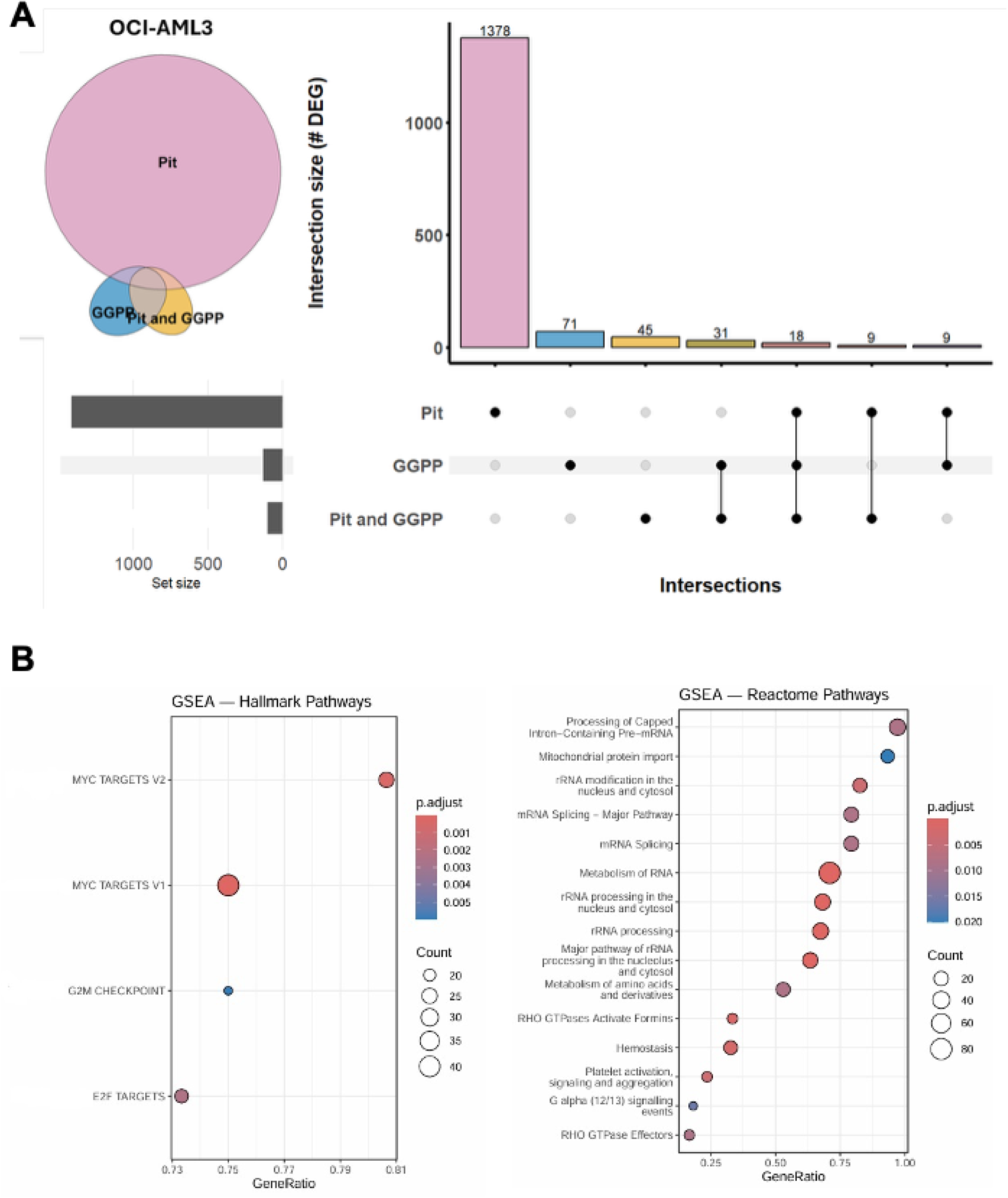
Transcriptome analysis identifies features of the response to PIT that depend on GGPP depletion. (A) Venn diagram and intersection plot showing DEGs in OCI-AML3 cells treated with PIT, GGPP, or PIT + GGPP, compared to vehicle. Note that in the presence of exogenous GGPP, PIT treatment induced very few gene expression changes. (B) GSEA for hallmark (left, including MYC and E2F targets) and reactome pathways (right, including mitochondrial protein import). The p value decreases from blue to red colors, and the diameter of each dot corresponds to gene set size.

An independent approach using Ingenuity Pathway Analysis software also identified MYC/E2F downregulation and PUMA upregulation, along with c-JUN upregulation, as cellular mechanisms induced by pitavastatin and GGTI-298 (Fig. S9D).

### Functional validation of transcription factor targets

To investigate the mechanism of reduced MYC target gene expression, we measured changes in MYC protein by western blot. Treatment of OCI-AML3 cells with pitavastatin reduced MYC expression, an effect that was reversed by mevalonate or GGPP and phenocopied by GGTI-298 and GGTI-2417, but not GGTI-2133 (Figure 5A, B; S12). Pitavastatin also reduced MYC protein amounts in primary AML samples UCI-01 and UCI-06 (Figure 5A). MYC protein downregulation was not observed in PL-21 cells (resistant to pitavastatin single-agent cytotoxicity) but was also minimal in HNT-34 cells (sensitive to pitavastatin cytotoxicity) (Figure 5A, B).

**Figure 5:**
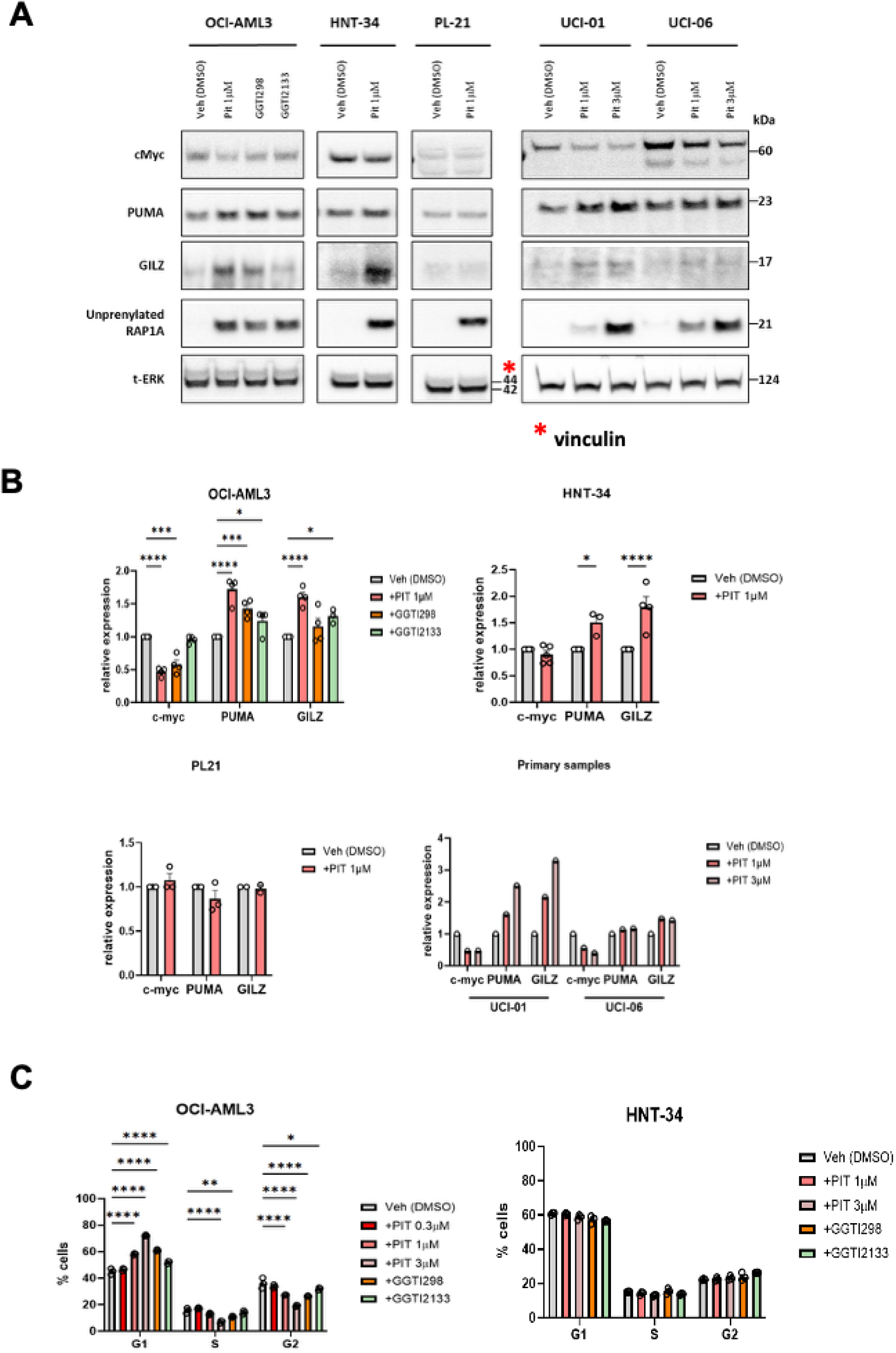
Pitavastatin treatment downregulates cMYC expression and increases PUMA and GILZ in AML cell lines and primary cells. (A) Two PIT-sensitive cell lines, one PIT-resistant cell line, and two PIT-sensitive primary AML samples were treated for 16hr as indicated and lysates subjected to western blotting with antibodies to cMYC, PUMA, GILZ, unprenylated RAP1A, and total ERK. (B) Graph of quantitation of western blots from 3-4 experiments for cell lines in panel A, or single experiment for two primary cell samples UCI-01 and UCI-06. Data are expressed as mean +/-SEM *p < 0.05, ***p < 0.001, ****p < 0.0001 two-way ANOVA with Dunnett’s test. (C) PIT and GGTI-298 cause G1 arrest in OCI-AML3 cells that are statin sensitive and show consistent MYC downregulation, but not in HNT-34 cells where MYC downregulation is less prominent. Data are expressed as mean +/-SEM *p < 0.05, ** p < 0.01, ****p < 0.0001 two-way ANOVA with Tukey’s test, n = 3.

Downregulation of MYC and E2F is commonly observed under conditions of cell cycle arrest (40,41). Accordingly, we found that pitavastatin and GGTI-298, but not GGTI-2133, increased the proportion of cells in G1 and decreased S phase in OCI-AML3 cells (Figure 5C). Pitavastatin and GGTI-298 had minimal effects on cell cycle in HNT-34 cells (Figure 5C) in alignment with the preservation of MYC protein expression (Figure 5A).

Pitavastatin strongly increased expression of the *TSC22D3* gene, with similar induction by GGTI-298 but to a lesser extent by GGTI-2133 (Fig. S10). *TSC22D3* encodes the glucocorticoid-induced leucine zipper (GILZ) protein that contributes to dexamethasone-induced apoptosis in MM cells (42). In accord with mRNA changes, protein measurements by western blot showed that pitavastatin increased GILZ expression in statin-sensitive cell lines but not in resistant PL-21 cells (Figure 5A, B). Addition of GGPP prevented the pitavastatin-induced GILZ expression (Fig. S13A). However, CRISPR/Cas9-mediated deletion of GILZ in OCI-AML3 cells did not affect cell viability changes in response to pitavastatin alone or the combination with venetoclax (Fig. S13B, C). These data suggest that GILZ induction is not essential for statin cytotoxicity.

### Pitavastatin suppresses oxidative metabolism and TCA cycle in OCI-AML3 cells

GSEA analysis showed that pitavastatin and/or GGTI-298 reduced gene sets associated with mitochondrial protein import and oxidative phosphorylation (Figure 3C, 4B). Combined analysis of data from OCI-AML3 cells in both transcriptome studies identified a set of 85 shared DEGs representing nuclear-encoded mitochondrial genes, of which only 3 were significantly changed by GGTI-2133 (Fig. 6A). A large majority of these were downregulated, including genes whose products regulate the function of components of the electronic transport chain (*CYCS*, *DHODH*, *NDUFAF4*, *TIMM21*, *TMEM126B*, *TMEM177*). The smaller subset of upregulated mitochondrial genes included *BBC3* (Figure 6A).

**Figure 6:**
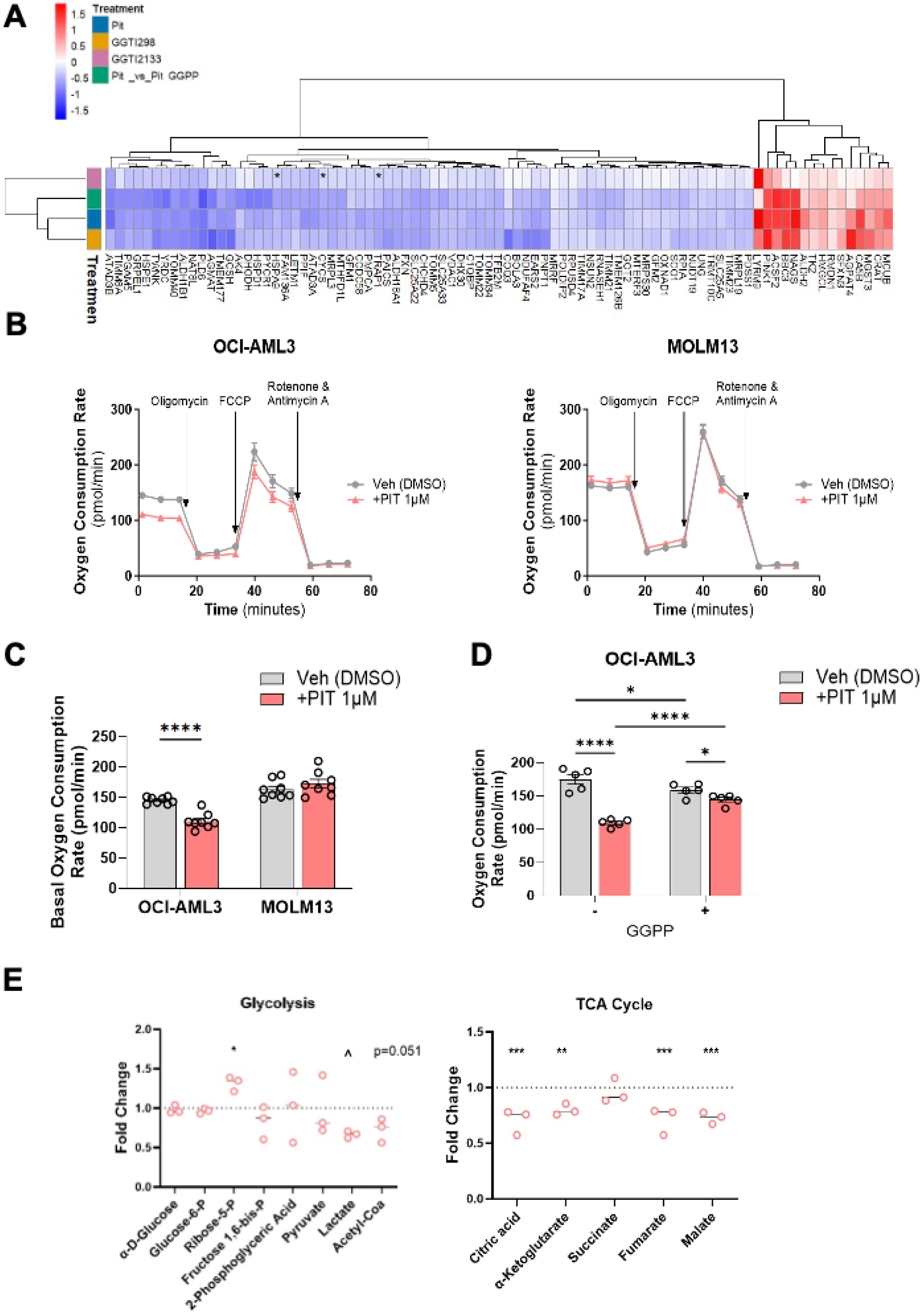
Functional assays provide evidence that PIT causes mitochondrial dysfunction. (A) Heat map of nuclear-encoded mitochondrial genes (MitoCarta database) across conditions in both transcriptome experiments. (B-D) Oxygen consumption rate (OCR) was measured by Seahorse metabolic flux analyzer. (B) Representative Seahorse assay traces in OCI-AML3 and MOLM13 cells treated with vehicle or pitavastatin for 16hr. (C) Graphs depict results of experiments as in panel B, shown as mean +/-SEM, **** p < 0.0001 two-way ANOVA with Sidak’s test, n = 8. (D) Effect of pitavastatin on basal OCR is significantly reversed by GGPP. * p < 0.05, **** p < 0.0001 two-way ANOVA with Fisher’s least squares difference test, n = 5. (E) ^13^C-glucose tracing identifies reduced TCA cycle intermediates in OCI-AML3 cells treated with pitavastatin for 16hr (13C-glucose added for last 2hr). Left graph shows quantitation of glycolytic metabolites, lactate, and acetyl-CoA, containing all ^13^C atoms traced. Right graph shows TCA cycle metabolites with ^13^C in M+2 position. Each dot plot depicts results of 3 biological replicates, each representing 4-6 technical replicates, normalized as a ratio of the pitavastatin-treated sample to the vehicle-treated group. * p < 0.05, ** p < 0.01, *** p < 0.001 two-way ANOVA with Fisher’s least squares difference test (^ p = 0.051 for lactate as marked).

The changes in expression of mRNAs encoding mitochondrial proteins prompted us to assess mitochondrial mass and function. Pitavastatin reduced expression of the mitochondrial protein TOM20 in OCI-AML3 cells but not MOLM13 cells (Fig. S14A); however, only one of two alternate assays for mitochondrial mass gave similar results (Fig. S14B, C). Seahorse metabolic flux analysis showed that pitavastatin reduced basal oxygen consumption rate (OCR) in OCI-AML3 cells but not in MOLM13 cells (Figure 6B, C), consistent with the lack of downregulation of mitochondrial genes in MOLM13. Addition of GGPP partially reversed the decreased basal OCR in OCI-AML3 cells treated with pitavastatin (Figure 6D). The combination of pitavastatin with venetoclax strongly and significantly reduced basal OCR and spare respiratory capacity in OCI-AML3 and MOLM13 cells (Fig. S15A, B). However, after 16hr of treatment with venetoclax +/-pitavastatin, a large percentage of cells had released cytochrome C (Fig. S15C, D), effects that might indirectly suppress OCR. Cytochrome C release did not occur with pitavastatin alone (Fig. S15C), a condition where basal OCR was reduced in OCI-AML3 cells (Figure 6B, C). Concurrent with decreased OCR, pitavastatin also caused loss of mitochondrial membrane potential in a significant percentage of OCI-AML3, as well as MOLM14 parental and *TP53* mutant lines (Fig. S16A, B).

To investigate whether mitochondrial dysfunction is associated with metabolic flux, we conducted untargeted metabolomics combined with [U-^13^C]-glucose tracing. The data revealed that, in OCI-AML3 cells, 16-hour pitavastatin treatment significantly reduced the production of TCA cycle intermediates from glucose (Figure 6E).

## Discussion

The combination of venetoclax with 5-azacitidine (VEN-AZA) is a standard of care regimen for AML patients who are >75 years old or unfit for intensive chemotherapy. As first-line therapy in previously untreated AML patients, VEN-AZA increases the frequency of complete remission and provides an increase in median OS compared to AZA alone (10). However, it has become clear that VEN-AZA provides no survival advantage compared to AZA alone in the subgroup with *TP53* loss of function (11,12). In addition, venetoclax resistance is common in AML cases with *FLT3*-ITD and/or *RAS* mutations, enhanced mitochondrial metabolism, or MCL-1 overexpression (8,13,14,18). Currently there are no clinically used drugs that can address multiple venetoclax resistance mechanisms. Here we show that an FDA-approved HMGCR inhibitor, pitavastatin, potentiates venetoclax cytotoxicity in multiple AML cell models including established and primary AML cells with *TP53* mutations or *FLT3*-ITD. In most statin-sensitive cell lines, pitavastatin upregulates the pro-apoptotic protein PUMA. In some cell lines, pitavastatin decreases MCL-1, and can inhibit oxidative metabolism.

The selective cytotoxicity of statins in AML compared to normal hematopoietic progenitors was first reported more than 30 years ago (43). Why has this not yet led to application in clinical practice? Three reasons deserve prominent attention. First, the great majority of cell-based assays in the literature employ concentrations of statins (5-20 µM) that are 100 – 1000 times higher than clinical achievable by the statin drugs most prescribed (i.e., simvastatin, atorvastatin, rosuvastatin). Thus, the cytotoxic effects and mechanisms described *in vitro* have questionable relevance to the clinical scenario. Second, it is essential to choose hydrophobic statin drugs with good bioavailability (30). Some prospective AML clinical trials have used hydrophilic statins (e.g., pravastatin; (44,45)) that, even at high doses, do not readily enter blood cells that lack active transport mechanisms. Third, statin pharmacology is different in rodents (46), resulting in inadequate inhibition of the mevalonate pathway and generally weak *in vivo* anti-cancer effects in preclinical models (33,47), which can diminish enthusiasm for advancing to human trials.

Pitavastatin has gained recognition as an ideal molecule for applications in oncology (30).

This compound is ∼3-times more potent on a molar basis in cell lines, achieves plasma concentrations in the 200 – 500 nM range in humans at the FDA-approved dose, and has an extended half-life and fewer drug interactions than other statins (21,26,48,49). Our phase 1 clinical trial found that daily 2 or 4 mg doses of pitavastatin were well tolerated in AML and CLL patients receiving venetoclax as standard of care (29). In this study, we show that pitavastatin is more potent than simvastatin in AML cell lines. While the concentration of pitavastatin that has single-agent cytotoxicity varies among AML cell lines from 0.3 to 3 µM, we observe activity in primary AML cells at concentrations in the 0.1 – 0.3 µM range. Importantly, pitavastatin is active in *TP53*-mutant samples regardless of their sensitivity to VEN or AZA. Pitavastatin enhanced venetoclax cytotoxicity in four AML cell lines with FLT3-ITD (MOLM13, MOLM14, MV4.11, PL-21) and in three out of five primary samples tested (UCI-03,-07,-09).

Selective inhibition of BCL2 with venetoclax is most effective in cancers that depend on BCL2 for survival. Basal or acquired dependence on MCL-1 is a common mechanism of venetoclax resistance in AML (34,50,51). Unfortunately, direct targeting of MCL-1 with highly selective compounds is associated with significant adverse events (52). Suppressing MCL-1 transcription via CDK9 inhibitors is another strategy under investigation (53). Pitavastatin is an alternative option, based on our observation that the compound can overcome venetoclax-induced increases in MCL-1 expression. In these experiments, we observed that in several AML cell lines, treatment with venetoclax or the combination of pitavastatin with venetoclax reduced PUMA protein amounts compared to basal. Thus, the mechanisms of cytotoxicity may differ in response to pitavastatin alone versus the combination with venetoclax.

There has been considerable interest in targeting oxidative metabolism to treat AML. Preclinical studies established that AML cell venetoclax resistance correlates with increased mitochondrial metabolism (17,18,54). Furthermore, various OXPHOS inhibitors potentiate cytotoxicity of venetoclax and prevent or overcome resistance (17,18,55,56). A selective inhibitor of complex-1 of the electron transport chain entered clinical trials but a phase I study reported significant toxicities and lack of efficacy at tolerated doses (57,58). Treatment with pitavastatin might provide an alternative, well-tolerated route to achieve a reduction in mitochondrial respiration. Our data show that at a time point before the onset of apoptosis, pitavastatin reduces basal OCR while triggering MMP loss in a significant fraction of cells. In OCI-AML3 cells, these mitochondrial defects were accompanied by decreased flux of glucose to the TCA cycle. In a separate paper, we reported that pitavastatin and rosuvastatin reverse the increase in OXPHOS observed in cytarabine-resistant *TP53*-mutant AML cells (33).

Mitochondrial dysfunction has the potential to reduce ATP production and cause activation of AMPK, a pathway that can promote PUMA expression via the integrated stress response (ISR) in AML cells (54). We previously showed that pitavastatin activates the ISR in MM cell lines; however, under the same conditions a chemical inhibitor of the ISR did not affect statin cytotoxicity or venetoclax sensitization in AML cell lines (26).

The effects of pitavastatin on AML cell survival, PUMA upregulation, and OXPHOS are dependent on GGPP depletion as they are largely rescued by exogenous GGPP. Furthermore, two inhibitors of GGTase-1 recapitulated many actions of pitavastatin, supporting the hypothesis that GGTase-1 substrates have essential functions in AML cell survival. A key question is how the mevalonate/GGPP axis programs AML transcription to promote proliferation and survival. To address this, we determined which transcriptional changes induced by pitavastatin are recapitulated by GGTI-298 and reversed by GGPP. Together these transcriptome analyses revealed several novel findings. First, the mevalonate pathway via GGPP promotes expression of cMYC and E2F target genes, correlating with cell cycle progression. Second, the mevalonate pathway regulates expression of many nuclear-encoded mitochondrial genes, some which are associated with OXPHOS. This connection might underly how treatment with pitavastatin reduces basal OCR and spare respiratory capacity and disrupts MMP. Notably, the extent of pitavastatin-induced transcriptional reprogramming is quite variable, with more prominent effects in OCI-AML3 cells compared to other lines tested here (Supplementary Excel files with volcano plots). This might be due in part to varying levels of positive feedback reactivation of the mevalonate pathway (59); indeed, pitavastatin-treated HNT-34 cells had a strong cholesterol pathway upregulation signature and overall fewer DEGs than OCI-AML3 cells. It is possible that the kinetics of transcriptional reprogramming differs among cell lines such that a shared DEG signature is difficult to define. Ongoing work aims to (i) identify transcription factors or epigenetic changes underlying altered gene expression and (ii) determine which geranylgeranylated protiens transmit key survival signals that are disrupted by statin treatment.

To broaden the utility of venetoclax in AML, it is essential to identify new combination regimens that address common mechanisms of resistance. In this study we have presented evidence that pitavastatin, an FDA-approved agent, has potent cytotoxic activity in human AML cells including samples with *TP53* loss. Pitavastatin treatment achieves several cellular changes that sensitize to venetoclax including PUMA upregulation, MCL-1 downregulation, and reduced respiration. Together with other findings from various laboratories (27,28,30), these results support further investigation of pitavastatin/venetoclax combinations in AML.

## Acknowledgements

The authors thank Dr. Elizabeth Brèm and Dr. Joel Leverson for helpful discussions. We acknowledge and appreciate research contributions from undergraduate students Nilufar Montazer, Ashley Navarro, Bita Sadrossadat, Jennifer Ly, Harrison Unger, and Enoc Ulloa. This work was supported by the UC Irvine Training Program for Interdisciplinary Cancer Research (T32-CA009054, to DJ), the UPenn Hematology Clinical Research Training Program (T32-HL0439, to SJS), research grants from the National Institutes of Health (R01-AA029124 to CJ; U54CA283759, to MC), an Impact Award from the Department of Defense Peer-Reviewed Cancer Research Program (W81XWH-20-1-0867, to DAF), a Veterans Merit Award (I01BX000918, to MC), a Discovery Boost grant from the American Cancer Society (DBG-24-1319247, to DAF), a Pew Scholar Award (to CJ), the American Society of Hematology (to SJS), American Society of Clinical Oncology (to SJS), and by an Anti-Cancer Challenge pilot grant (to DAF) from the Chao Family Comprehensive Cancer Center (supported by P30-CA062203). The project used Shared Resources of the Chao Family Comprehensive Cancer Center (Optical Biology Core; Biostatistics Shared Resource; Experimental Tissue Resource).

## Author Contributions

Supervision: RB, DJ, MP, SJS, CJ, MC, DAF. Funding acquisition: SJS, SMS, CJ, MC, DAF. Conceptualization: RB, DJ, MP, SJS, MC, DAF. Methodology/Provision of models/reagents: SJS, GW, SMS, MK, AGF, CJ, MC. Data collection: RB, DJ, MP, SJS, IBW, IT, ZY, IL, AB, MK. Data analysis and interpretation: RB, DJ, MP, SJS, IBW, IT, ZY, IL, CJ, MC, DAF. Bioinformatics: RB, DJ, MP. Statistical analysis: RB, DJ, MP, SJS. Figure preparation: RB. Writing – original draft: RB, DJ, MP, DAF. Critical review of the manuscript & writing – review and editing prior to finalization: All authors.

## Data Availability

RNA-seq datasets produced in this study will be available in the GEO databases (deposit is in progress).

---

**Table S2.**

Characteristics of primary AML samples
| Sample | Age | Sex | Disease | Disease Status | Molecular Cytogenetics | Treatment History |
| --- | --- | --- | --- | --- | --- | --- |
| UCI-01 | 78 | M | AML | New Diagnosis | 46,XY,+19(3)/46,XY,add(19)(P13)(3)/46,XY(14) | No prior treatment |
| UCI-02 | 76 | M | AML arising from MDS | New Diagnosis | 44-45, XY, add (1) (p36.3), add (3) (P13), -4, add(5)(q11.2),-7 (q22),+8del(9) (q32),-10, add(17) (P11.2),-20,+2-3mar(cp16) | Azacitidine followed by Decitabine for MDS prior to AML diagnosis |
| UCI-03 | 59 | F | AML | Relapse | FLT3 ITD, NPM1 XX | Idarubicin/Cytarabine. |
| UCI-04 | 61 | F | AML M5 | New Diagnosis | NPM1 mutant XX | No prior treatment |
| UCI-06 | 26 | F | AML | New Diagnosis | CEBPA mutant (p.Asp301_Leu315dup)<br>KIT mutant (p.Asp816His;) XX | No prior treatment |
| UCI-07 | 74 | M | AML | New Diagnosis | NPM1 mutant (p.Trp288CysfsTer12)<br>FLT3-ITD (ITD Size: 107bp) XY | No prior treatment |
| UCI-08 | 50 | F | AML | New Diagnosis | NPM1 (p.Trp288CysfsTer12)<br>DNMT3A mutant (p.Arg882His) XX | No prior treatment |
| UCI-09 | 83 | F | AML | New Diagnosis | NPM1 (p.Trp288CysfsTer12)<br>FLT3-ITD (ITD Size: 26pb) XX | No prior treatment |
| UCI-10 | 41 | F | AML | New Diagnosis | FLT3-ITD | No prior treatment |
| UCI-11 | 77 | M | MDS/MPN transformed to AML | Transformation to AML | complex karyotype, monosomy 7, deletion of 12p,and isochromosome 17q | Hydrea for MDS/MPN prior to AML diagnosis |
| UCI-12 | 43 | F | AML | Relapse | FLT3-ITD | Idarubicin/Cytarabine |
| LBV-03 | 80 | M | AML | New Diagnosis | Trisomy 8 / duplication of 8q22 and gain of KMT2A | No prior treatment |
| UPENN-01 |  |  | AML | New Diagnosis | TP53 frameshift mutation with der(17)t(11;17) |  |
| TUPENN-02 |  |  | AML | New Diagnosis | TP53 missense Y220C, CBL, NPM1, TET2 mutations, VUS EGR2 |  |

## Notes

### Summary of Updates

The full title and abstract were inadvertently shortened during the transfer process from the journal where we originally submitted the manuscript. The revisions restore the original title and abstract.

## References

1. Cory S, Roberts AW, Colman PM, Adams JM. Targeting BCL-2-like Proteins to Kill Cancer Cells. Trends Cancer. 2016 Aug;2(8):443–60.

2. Leverson JD, Sampath D, Souers AJ, Rosenberg SH, Fairbrother WJ, Amiot M, et al. Found in Translation: How Preclinical Research Is Guiding the Clinical Development of the BCL2-Selective Inhibitor Venetoclax. Cancer Discov. 2017 Dec;7(12):1376–93.

3. Merino D, Kelly GL, Lessene G, Wei AH, Roberts AW, Strasser A. BH3-Mimetic Drugs: Blazing the Trail for New Cancer Medicines. Cancer Cell. 2018 Dec 10;34(6):879–91.

4. Roberts AW, Davids MS, Pagel JM, Kahl BS, Puvvada SD, Gerecitano JF, et al. Targeting BCL2 with Venetoclax in Relapsed Chronic Lymphocytic Leukemia. N Engl J Med. 2016 Jan 28;374(4):311–22.

5. DiNardo CD, Pratz K, Pullarkat V, Jonas BA, Arellano M, Becker PS, et al. Venetoclax combined with decitabine or azacitidine in treatment-naive, elderly patients with acute myeloid leukemia. Blood. 2019 Jan 3;133(1):7–17.

6. Grant S. Rational combination strategies to enhance venetoclax activity and overcome resistance in hematologic malignancies. Leuk Lymphoma. 2017 Aug 24;59(6):1–8.

7. Nechiporuk T, Kurtz SE, Nikolova O, Liu T, Jones CL, D’Alessandro A, et al. The TP53 apoptotic network is a primary mediator of resistance to BCL2 inhibition in AML cells. Cancer Discov. 2019 Jul;9(7):910–25.

8. DiNardo CD, Tiong IS, Quaglieri A, MacRaild S, Loghavi S, Brown FC, et al. Molecular patterns of response and treatment failure after frontline venetoclax combinations in older patients with AML. Blood. 2020 Mar 12;135(11):791–803.

9. Thijssen R, Diepstraten ST, Moujalled D, Chew E, Flensburg C, Shi MX, et al. Intact TP-53 function is essential for sustaining durable responses to BH3-mimetic drugs in leukemias. Blood. 2021 May 20;137(20):2721–35.

10. DiNardo CD, Jonas BA, Pullarkat V, Thirman MJ, Garcia JS, Wei AH, et al. Azacitidine and venetoclax in previously untreated acute myeloid leukemia. N Engl J Med. 2020 Aug 13;383(7):617–29.

11. Pollyea DA, Pratz KW, Wei AH, Pullarkat V, Jonas BA, Recher C, et al. Outcomes in Patients with Poor-Risk Cytogenetics with or without TP53 Mutations Treated with Venetoclax and Azacitidine. Clin Cancer Res. 2022 Dec 15;28(24):5272–9.

12. Döhner H, Pratz KW, DiNardo CD, Wei AH, Jonas BA, Pullarkat VA, et al. Genetic risk stratification and outcomes among treatment-naive patients with AML treated with venetoclax and azacitidine. Blood. 2024 Nov 21;144(21):2211–22.

13. Bhatt S, Pioso MS, Olesinski EA, Yilma B, Ryan JA, Mashaka T, et al. Reduced mitochondrial apoptotic priming drives resistance to BH3 mimetics in acute myeloid leukemia. Cancer Cell. 2020 Dec 14;38(6):872–890.e6.

14. Sango J, Carcamo S, Sirenko M, Maiti A, Mansour H, Ulukaya G, et al. RAS-mutant leukaemia stem cells drive clinical resistance to venetoclax. Nature. 2024 Dec;636(8041):241–50.

15. Brown FC, Wang X, Birkinshaw R, Chua CC, Morley T, Kasapgil S, et al. Acquired BCL2 variants associated with venetoclax resistance in acute myeloid leukemia. Blood Adv. 2025 Jan 14;9(1):127–31.

16. Sharon D, Cathelin S, Mirali S, Di Trani JM, Yanofsky DJ, Keon KA, et al. Inhibition of mitochondrial translation overcomes venetoclax resistance in AML through activation of the integrated stress response. Sci Transl Med. 2019 Oct 30;11(516).

17. Liu F, Kalpage HA, Wang D, Edwards H, Hüttemann M, Ma J, et al. Cotargeting of Mitochondrial Complex I and Bcl-2 Shows Antileukemic Activity against Acute Myeloid Leukemia Cells Reliant on Oxidative Phosphorylation. Cancers (Basel). 2020 Aug 24;12(9).

18. Bosc C, Saland E, Bousard A, Gadaud N, Sabatier M, Cognet G, et al. Mitochondrial inhibitors circumvent adaptive resistance to venetoclax and cytarabine combination therapy in acute myeloid leukemia. Nat Cancer. 2021 Nov 11;2(11):1204–23.

19. Glytsou C, Chen X, Zacharioudakis E, Al-Santli W, Zhou H, Nadorp B, et al. Mitophagy promotes resistance to BH3 mimetics in acute myeloid leukemia. Cancer Discov. 2023 Jul 7;13(7):1656–77.

20. Mullen PJ, Yu R, Longo J, Archer MC, Penn LZ. The interplay between cell signalling and the mevalonate pathway in cancer. Nat Rev Cancer. 2016 Nov;16(11):718–31.

21. Juarez D, Fruman DA. Targeting the mevalonate pathway in cancer. Trends Cancer. 2021 Jun;7(6):525–40.

22. Clendening JW, Penn LZ. Targeting tumor cell metabolism with statins. Oncogene. 2012 Nov 29;31(48):4967–78.

23. Longo J, van Leeuwen JE, Elbaz M, Branchard E, Penn LZ. Statins as anticancer agents in the era of precision medicine. Clin Cancer Res. 2020 Nov 15;26(22):5791–800.

24. Tripathi S, Gupta E, Galande S. Statins as anti-tumor agents: A paradigm for repurposed drugs. Cancer Rep (Hoboken). 2024 May;7(5):e2078.

25. Lee JS, Roberts A, Juarez D, Vo T-TT, Bhatt S, Herzog L-O, et al. Statins enhance efficacy of venetoclax in blood cancers. Sci Transl Med. 2018 Jun 13;10(445).

26. Juarez D, Buono R, Matulis SM, Gupta VA, Duong M, Yudiono J, et al. Statin-induced Mitochondrial Priming Sensitizes Multiple Myeloma Cells to BCL2 and MCL-1 Inhibitors. Cancer Res Commun. 2023 Dec 8;3(12):2497–509.

27. Gimenez N, Tripathi R, Giró A, Rosich L, López-Guerra M, López-Oreja I, et al. Systems biology drug screening identifies statins as enhancers of current therapies in chronic lymphocytic leukemia. Sci Rep. 2020 Dec 17;10(1):22153.

28. Al-Zebeeby A, Vogler M, Milani M, Richards C, Alotibi A, Greaves G, et al. Targeting intermediary metabolism enhances the efficacy of BH3 mimetic therapy in hematologic malignancies. Haematologica. 2019;104(5):1016–25.

29. Brem EA, Shieh K, Juarez D, Buono R, Jeyakumar D, O’Brien S, et al. A phase 1 study of adding pitavastatin to venetoclax-based therapy in AML and CLL/SLL: a tolerable, mechanism-based, drug repurposing strategy. Blood Neoplasia. 2024 Aug;100036.

30. Abdullah MI, de Wolf E, Jawad MJ, Richardson A. The poor design of clinical trials of statins in oncology may explain their failure - Lessons for drug repurposing. Cancer Treat Rev. 2018 Sep;69:84–9.

31. Wang M, Casey PJ. Protein prenylation: unique fats make their mark on biology. Nat Rev Mol Cell Biol. 2016 Feb;17(2):110–22.

32. Han L, Zhang Q, Dail M, Shi C, Cavazos A, Ruvolo VR, et al. Concomitant targeting of BCL2 with venetoclax and MAPK signaling with cobimetinib in acute myeloid leukemia models. Haematologica. 2020 Mar;105(3):697–707.

33. Skuli SJ, Bakayoko A, Kruidenier M, Manning B, Pammer P, Salimov A, et al. Chemoresistance of TP53 mutant acute myeloid leukemia requires the mevalonate byproduct, geranylgeranyl pyrophosphate, for induction of an adaptive stress response. Leukemia. 2025 Jul 9;39:2087–2098.

34. Teh TC, Nguyen N-Y, Moujalled DM, Segal D, Pomilio G, Rijal S, et al. Enhancing venetoclax activity in acute myeloid leukemia by co-targeting MCL1. Leukemia. 2017 Jul 28;32(2):303–12.

35. Caenepeel S, Brown SP, Belmontes B, Moody G, Keegan KS, Chui D, et al. AMG 176, a Selective MCL1 Inhibitor, Is Effective in Hematologic Cancer Models Alone and in Combination with Established Therapies. Cancer Discov. 2018 Dec;8(12):1582–97.

36. Xia Z, Tan MM, Wong WW, Dimitroulakos J, Minden MD, Penn LZ. Blocking protein geranylgeranylation is essential for lovastatin-induced apoptosis of human acute myeloid leukemia cells. Leukemia. 2001 Sep;15(9):1398–407.

37. Wong WW-L, Clendening JW, Martirosyan A, Boutros PC, Bros C, Khosravi F, et al. Determinants of sensitivity to lovastatin-induced apoptosis in multiple myeloma. Mol Cancer Ther. 2007 Jun;6(6):1886–97.

38. Xia Y, Xie Y, Yu Z, Xiao H, Jiang G, Zhou X, et al. The mevalonate pathway is a druggable target for vaccine adjuvant discovery. Cell. 2018 Nov 1;175(4):1059–1073.e21.

39. Peng H, Carrico D, Thai V, Blaskovich M, Bucher C, Pusateri EE, et al. Synthesis and evaluation of potent, highly-selective, 3-aryl-piperazinone inhibitors of protein geranylgeranyltransferase-I. Org Biomol Chem. 2006 May 7;4(9):1768–84.

40. Schuldiner O, Benvenisty N. A DNA microarray screen for genes involved in c-MYC and N-MYC oncogenesis in human tumors. Oncogene. 2001 Aug 16;20(36):4984–94.

41. Baudino TA, Maclean KH, Brennan J, Parganas E, Yang C, Aslanian A, et al. Myc-mediated proliferation and lymphomagenesis, but not apoptosis, are compromised by E2f1 loss. Mol Cell. 2003 Apr;11(4):905–14.

42. Kervoëlen C, Ménoret E, Gomez-Bougie P, Bataille R, Godon C, Marionneau-Lambot S, et al. Dexamethasone-induced cell death is restricted to specific molecular subgroups of multiple myeloma. Oncotarget. 2015 Sep 29;6(29):26922–34.

43. Newman A, Clutterbuck RD, Powles RL, Millar JL. Selective inhibition of primary acute myeloid leukaemia cell growth by simvastatin. Leukemia. 1994 Nov;8(11):2023–9.

44. Kornblau SM, Banker DE, Stirewalt D, Shen D, Lemker E, Verstovsek S, et al. Blockade of adaptive defensive changes in cholesterol uptake and synthesis in AML by the addition of pravastatin to idarubicin + high-dose Ara-C: a phase 1 study. Blood. 2007 Apr 1;109(7):2999–3006.

45. Advani AS, Li H, Michaelis LC, Medeiros BC, Liedtke M, List AF, et al. Report of the relapsed/refractory cohort of SWOG S0919: A phase 2 study of idarubicin and cytarabine in combination with pravastatin for acute myelogenous leukemia (AML). Leuk Res. 2018 Apr;67:17–20.

46. Schonewille M, de Boer JF, Mele L, Wolters H, Bloks VW, Wolters JC, et al. Statins increase hepatic cholesterol synthesis and stimulate fecal cholesterol elimination in mice. J Lipid Res. 2016 Aug;57(8):1455–64.

47. Hillis AL, Martin TD, Manchester HE, Högström J, Zhang N, Lecky E, et al. Targeting Cholesterol Biosynthesis with Statins Synergizes with AKT Inhibitors in Triple-Negative Breast Cancer. Cancer Res. 2024 Oct 1;84(19):3250–66.

48. Catapano AL. Pitavastatin - pharmacological profile from early phase studies. Atheroscler Suppl. 2010 Dec;11(3):3–7.

49. Catapano AL. Pitavastatin: a different pharmacological profile. Clin Lipidol. 2012 Jun;7(3s):3–9.

50. Moujalled DM, Pomilio G, Ghiurau C, Ivey A, Salmon J, Rijal S, et al. Combining BH3-mimetics to target both BCL-2 and MCL1 has potent activity in pre-clinical models of acute myeloid leukemia. Leukemia. 2019;33(4):905–17.

51. Zhang Q, Riley-Gillis B, Han L, Jia Y, Lodi A, Zhang H, et al. Activation of RAS/MAPK pathway confers MCL-1 mediated acquired resistance to BCL-2 inhibitor venetoclax in acute myeloid leukemia. Signal Transduct Target Ther. 2022 Feb 21;7(1):51.

52. Desai P, Lonial S, Cashen A, Kamdar M, Flinn I, O’Brien S, et al. A Phase 1 First-in-Human Study of the MCL-1 Inhibitor AZD5991 in Patients with Relapsed/Refractory Hematologic Malignancies. Clin Cancer Res. 2024 Nov 1;30(21):4844–55.

53. Davids MS, Brander DM, Alvarado-Valero Y, Diefenbach CS, Egan DN, Dinner SN, et al. A phase 1 study of the CDK9 inhibitor voruciclib in relapsed/refractory acute myeloid leukemia and B-cell malignancies. Blood Adv. 2025 Feb 25;9(4):820–32.

54. Grenier A, Poulain L, Mondesir J, Jacquel A, Bosc C, Stuani L, et al. AMPK-PERK axis represses oxidative metabolism and enhances apoptotic priming of mitochondria in acute myeloid leukemia. Cell Rep. 2022 Jan 4;38(1):110197.

55. Hua L, Yang N, Li Y, Huang K, Jiang X, Liu F, et al. Metformin sensitizes AML cells to venetoclax through endoplasmic reticulum stress-CHOP pathway. Br J Haematol. 2023 Jul 6;202(5):971–84.

56. Velez J, Pan R, Lee JTC, Enciso L, Suarez M, Duque JE, et al. Biguanides sensitize leukemia cells to ABT-737-induced apoptosis by inhibiting mitochondrial electron transport. Oncotarget. 2016 Jun 6;7(32):51435–49.

57. Molina JR, Sun Y, Protopopova M, Gera S, Bandi M, Bristow C, et al. An inhibitor of oxidative phosphorylation exploits cancer vulnerability. Nat Med. 2018 Jul;24(7):1036– 46.

58. Yap TA, Daver N, Mahendra M, Zhang J, Kamiya-Matsuoka C, Meric-Bernstam F, et al. Complex I inhibitor of oxidative phosphorylation in advanced solid tumors and acute myeloid leukemia: phase I trials. Nat Med. 2023 Jan 19;29(1):115–26.

59. Pandyra A, Mullen PJ, Kalkat M, Yu R, Pong JT, Li Z, et al. Immediate utility of two approved agents to target both the metabolic mevalonate pathway and its restorative feedback loop. Cancer Res. 2014 Sep 1;74(17):4772–82.

