## Supplementary Materials for "Pitavastatin counteracts venetoclax resistance mechanisms in acute myeloid leukemia by depleting geranylgeranyl pyrophosphate"

**Supplementary Materials and Methods**

Chemicals

Pitavastatin (Fisher Scientific #NC0110176), venetoclax (Selleck Chemicals #S8048), S63845 (Selleck S8383/Chemgood C-1370), GGTI-298 (Fisher Scientific # HY-15871), GGTI-2133 (Sigma Aldrich #G5294), THZ145 (MedChem Express #HY-150168) and GGTI2417 (synthesized as described (1)) were dissolved in DMSO. Mevalonate (Sigma-Aldrich #90469) was resuspended with a 7:3 solution of methanol and 10 mM ammonium hydroxide. GGPP (Sigma #G6025-1VL or Cayman Chemical 63330) and FPP (Cayman Chemical #63250) were used at 5 μg/ml.

Cell Culture

Cell lines were maintained at concentrations between 0.5 million to 2 million per ml. Experiments requiring incubation times less than 24 hours were plated at 1 million per ml while greater than 24 hours were plated at 0.5 million per ml. MOLM14 parental cells and *TP53* mutant were cultured in alphaMEM supplemented with 10% FBS.

Cell Viability

Cell viability assays were performed in 96-well format for 48 hours. $5 \times 10^4$ cells were cultured in 200 μL growth medium with inhibitors for 48 hours. Inhibitors did not exceed a 0.1% DMSO concentration. Viability was determined by fluorescence of propidium iodide (Life Technologies #P3566) and/or Annexin V Alexa Fluor 647 conjugate (Life Technologies #A23204) measured by the Novocyte 3000 (Agilent) or Attune NxT (Thermo Fisher). Viability of cells was quantified using FlowJo software v10.10.0 (FlowJo LLC).

Cell Cycle Analysis

After 48h treatment cells were harvested and washed in PBS with 5 mM EDTA, then fixed in cold ethanol for 30min at room temperature. Cells were spun down, washed and treated with ribonuclease for 30min at 37°C. Cells were stained with propidium iodide, and 10,000 cellular events were collected by flow cytometry using Attune NxT (ThermoFisher Scientific). Cell cycle was quantified using FlowJo software v10.10.0 (FlowJo LLC).

Western Blotting

Cells after treatment were lysed in radio-immunoprecipitation assay (RIPA) buffer (150 mM NaCl, 1.0% IGEPAL CA- 630, 0.5% sodium deoxycholate, 0.1% SDS, and 50 mM Tris, pH 8.0, 2 mM EDTA, 50 mM NaF) supplemented with protease inhibitor cocktail (Sigma-Aldrich #539134-1ML) and phosphatase inhibitor cocktails 2 and 3 (Sigma-Aldrich #P5726-1ML #P0044-5ML). Protein concentrations were normalized using a Bradford protein assay (Bio-Rad #5000006). Lysates were prepared at 1 µg/µl concentration in 1X XT Sample Buffer (Bio-Rad #1610791) and 5% 2- mercaptoethanol (Sigma-Aldrich). Lysates were run on 4-12% Bolt Bis-Tris Plus gels (Life Technologies) and transferred onto nitrocellulose membranes.

The filters were blocked in 5% milk for 1h at room temperature and then incubated overnight at 4° with specific antibodies: PUMA (1:1000 Cell Signaling #12450), unprenylated RAP1A (1:50 Santa Cruz sc-373968), p44/42 MAPK (Erk1/2) (1:1000 Cell Signaling #4696), c-Myc (1:1000 Cell Signaling #18583) Vinculin (1:1000 Cell Signaling #13901), GILZ (1:1000 ThermoFisher Scientific # 12352-1-AP), Mcl1 (1:1000 Cell Signaling #39224). The following secondary HRP-conjugated antibodies were used: anti-mouse IgG, anti-rabbit IgG (1:5000 Promega #W4021 #W4011).

Chemiluminescence detected using a Nikon D700 SLR camera or the SynGene G:Box. Images were processed with auto-contrast uniformly across the entire image and densitometry was performed using ImageJ software.

Plasmid cloning, virus production and infection

CRISPR-Cas9–mediated gene knockout was generated using the lentiCRISPR v2 vector (Addgene plasmid #52961). For sgRNA cloning, the vector was linearized with BsmBI (New England Biolabs) and ligated with gene-specific sgRNAs using T4 DNA ligase (New England Biolabs). Guide RNAs targeting *BBC3* (PUMA; CGCTGGGCACGGGCGACTCC) or *TSC22D3* (GILZ; GGUGUUCUCACGCUCUAGCU) were cloned into the vector, and all constructs were confirmed by Sanger sequencing. Lentiviral particles were produced by co-transfecting 293T HEK cells with the sgRNA containing lentiCRISPR v2 plasmid (Addgene plasmid #52961) and the packaging plasmids pCMV-VSVG (Addgene plasmid #454) and psPAX2 (Addgene plasmid #12260), using TransIT-LT1 Transfection Reagent (Mirus Bio #MIR6703) according to the manufacturer's instructions. Viral supernatant was collected 72 hours post-transfection. Target cells ($1 \times 10^5$) were infected with a virus-containing supernatant and incubated at 37°C with 5% $CO_2$ for 5-7 days. Transduced cells were selected with puromycin at 2 µg/mL for 5–7 days, followed by 1 µg/mL for an additional 5 days, and maintained in 0.5 µg/mL thereafter. Gene knockout was confirmed by Western blotting in each experiment. Single-cell clones were generated by seeding transduced cells into 96-well plates at a density of 0.3 cells/well. Clonal populations were expanded, and those with confirmed gene knockout by western blotting were selected for downstream assays.

RNA Sequencing and Analysis

Cells were treated for 16 hours with inhibitors before being pelleted, washed, and flash frozen prior to RNA extraction. RNA was extracted using RNeasy Mini kit (QIAGEN, Germantown, MD, USA #74104) or Direct-zol RNA Microprep Kit (Thomas Scientific #1159U97) followed by on-column DNase I digestion as per manufacturer's instructions. The purity and concentration of the extracted RNA were assessed using Nanodrop before downstream applications. RNA was sent to UCI Genomic High Throughput Facility for library preparation and sequencing. In brief, library preparation was conducted using the TruSeq Stranded mRNA kit. Multiplexed samples were sequenced on the Illumina HiSeq 4000. A minimum of 40 million paired end reads were acquired for differential expression analysis and differential transcript utilization analysis. Raw fastq files generated by sequencing were quality assessed by Fastqc/0.11.9. Transcript-level quantification was performed on a high-performance compute cluster using Salmon v1.8.0 in quasi-mapping mode and a pre-built transcriptome index obtained from the DataBio RefGenomes repository (hg38/salmon_sa_index) made from the GCA_000001405.15 GRCh38_no_alt_analysis_set from NCBI. Transcript quantification for paired-end RNA-seq libraries was performed using the –validateMappings, --seqBias, and –gcBias flags to enable selective alignment and bias correction. –useVBOpt invoked variational Bayesian optimization and –numBootstraps 30 generated 30 bootstrap replicates per sample for transcript level to gene level conversion. Outputs were imported with tximeta using countsFromAbundance= “scaledTPM” for length-scaled counts. Transcript abundances were summarized to gene level with summarizeToGene. Gene-level differential expression analysis used DESeq2 with ~ batch + condition design. Batch effects were removed with limma removeBatchEffect and matrix was rlog transformed for sample-distance heatmaps. Shrunken log2fold changes were obtained with apeglm for visualizations. Significance threshold was padj <0.05. To compare DEGs across treatments, we generated two complementary visual summaries: UpSet plots using Complex Upset package in R and scaled ellipse Venn diagrams using eulerr. We also assessed DEGs by preranked GSEA with clusterProfiler using rank= sign(log2FC)*-log10(padj) from DESeq2 output as the rank metric per treatment or the average of rank per of two treatments within a cell line. We queried Reactome via Reactome PA, and Hallmark via msigdbr. Significant GSEA results were visualized by enrichplot dot plots. These significant GSEA results were compared across groups (either treatments within a cell line or the same treatment across cell lines) in R with terms filtered at q≤ 0.05 in dot plots built with ggplot2 with shared terms being included and outlined along with top 20 terms per group. Heatmaps were made in pheatmap using normalized counts from DESeq2. Counts were transformed as log2(count+1) and averaged per gene within each cell line and treatment group. For each gene, per-cell-line log2 fold changes were calculated as the difference between the mean log2 expression of each treatment and the corresponding vehicle control. To facilitate visual comparison of induction and repression patterns of significant genes across conditions in different genetic backgrounds, log2 fold changes were range-scaled within each cell line (and as such within each generalized linear model) so that the log2 fold change of gene each across all treatments is divided by the largest absolute log2 fold change for that gene in that line at the expense of absolute magnitude. These were plotted with hierarchical clustering of both genes and treatment conditions (Euclidean distance, complete linkage) to group genes by similarity in directional pattern. Absolute differences were visualized by bar plots in ggplot2 of DESeq2 apeglm shrunken log 2 fold changes and their standard errors (lfcSE).

Seahorse Assay

XF assays were performed using the Agilent Seahorse XFe96 Extracellular Flux Analyzer. The day before the assay, the sensor cartridge was placed into the calibration buffer medium supplied by Seahorse Biosciences to hydrate overnight. Seahorse XFe96 microplates wells were coated with 25 µL of Cell-Tak (Corning; Cat#354240) solution at a concentration of 22.4 µg/ml at room temperature for 20 minutes, washed twice with distilled water, and kept at 4 °C overnight. On the day of the experiment, AML cells were plated at a density of 80,000 cells per well for cell lines in XF base minimal DMEM media containing 11 mM glucose, 1 mM pyruvate and 2 mM glutamine. Then 180 µL of XF base minimal DMEM medium was added to each well and the microplate was centrifuged at 100 x g for 1 min with no break. After no more than one hour of incubation at 37 °C in CO2 free-atmosphere, basal oxygen consumption rate (OCR, as a mitochondrial respiration indicator) and extracellular acidification rate (ECAR, as a glycolysis indicator) were performed using the Seahorse XF Cell Mito Stress Test Kit (#103015-100).

Mitochondrial Membrane Potential

Cells were treated for 16 hr with inhibitors before a subset of untreated cells wee stained with 30 µM carbonyl cyanide m-chlorophenylhydrazone (CCCP) (MedChemExpress #HY-100941) for 15 min, away from light at 37°C, 5% $CO_2$. Cells were then stained with 100 nM tetramethyl-rhodamine ethyl ester perchlorate (TMRE) (MedChemExpress #HY-D0985A) for 15 min, away from light at 37°C, 5% $CO_2$. 10,000 events were collected by an Agilent Novocyte Quanteon flow cytometer. Percent mitochondrial membrane potential loss was determined on FlowJo by sub-gating on live cells in the EYFP channel that have shifted relative to the vehicle cells.

Measurement of mitochondrial mass

Cells were treated for 16 hours with inhibitors before a subset of untreated cells are stained with 30 µM CCCP (MedChemExpress #HY-100941) for 15 minutes, away from light at 37°C, 5% $CO_2$. Cells were then stained for cardiolipin with 250 nM Acridine Orange 10-Nonyl Bromide (NAO) (MedChemExpress #HY-D0993) and Zombie NIR Fixable Viability kit (total dilution 1:10,000) (Biolegend #423105) for 15 minutes, away from light at 37°C, 5% $CO_2$. 50,000 events were collected by an Agilent Novocyte Quanteon flow cytometer. MFI was determined by first gating on live cells using the APC/Cy7 channel, then sub-gated on cells using the Pacific Orange channel. MFI was then calculated from the histogram curve.

Mitochondrial mass in viable cells was measured by flow cytometry using Tom20 Antibody (5-10) Alexa Fluor488 (Santa Cruz #17764)

Mitochondrial DNA copy number

Cells were treated for 16 hours with inhibitors before being pelleted and lysed with mouse genotyping lysis buffer and Proteinase K (ThermoFisher Scientific #EO0491). Genomic DNA was isolated using phenol-chloroform 1:1 extraction, and ethanol precipitated overnight. DNA was spun down at maximum speed for 30 minutes, washed with 70% ethanol, and re-spun at maximum speed for another 30 minutes to remove remaining ethanol. The DNA pellet was resuspended in TE buffer. Quantitative PCR was run on each condition in triplicates with genomic DNA primers for *β2M* (forward: CGACGGGAGGGTCGGGACAA; reverse: GCCCCGCGAAAGAGCGGAAG) or mitochondrial DNA primers for *ND1* (forward: GTCAACCTCGCTTCCCCACCCT; reverse: TCCTGCGAATAGGCTTCCGGCT), and PowerUp SYBR Green Master Mix (ThermoFisher Scientific #A25742). Mitochondrial DNA copy number was calculated using the following equation:

$\Delta C_t$ = *ND1* average $C_t$ - *β2M* average $C_t$

Mitochondrial DNA copy number = $(1/(2^{\Delta Ct})) * 2$

Cytochrome C staining

Cells after treatment were washed in PBS and resuspended in Newmeyer Buffer. Cells were permeabilized with 0.01% digitonin for 5 min, fixed in 4% PFA for 20 min. After neutralizing PFA with neutralizing buffer (1.7M Tris Base, 1.25M Glycine, pH 9.1) samples were stained overnight with Alexa Fluor 647 Cytocrome C (#612310 Biolegend) in 10X tween intracellular staining buffer. Cytochrome c release was measured by the Novocyte 3000 or Novocyte Quanteon (Agilent) and quantified using FlowJo software v10.10.0 (FlowJo LLC).

Metabolomics

Cells were harvested and snap-frozen on dry ice. An ice-cold mixture of methanol:acetonitrile:water (40:40:20, vol; 0.5-0.6 mL) was added to samples and centrifuged at

16,000 x g for 10 min at 4°C. Supernatants (3 μL) from samples were analyzed as described (2). Briefly, a quadrupole-orbitrap mass spectrometer (Q Exactive Plus, ThermoFisher Scientific) operated in negative or positive ionization mode was coupled to a Vanquish Ultra High-Performance LC system (Thermo Fisher Scientific) with electrospray ionization. Scan range was m/z 70-1000, scanning frequency was 2 Hz and resolution was 140,000. LC separations were conducted using a XBridge BEH Amide column (2.1 mm x 150 mm2, 2.5 μm particle size, 130Å pore size pore size; Waters Corporation) with a gradient consisting of solvent A (20 mM ammonium acetate, 20 mM ammonium hydroxide in 95:5 water:acetonitrile, pH 9.45) and solvent B (acetonitrile). The flow rate was 0.150 mL/min. The gradient was: 0 min,85% B; 2 min, 85% B; 3 min, 80% B; 5 min, 80% B; 6 min, 75% B; 7 min, 75% B; 8 min, 70% B; 9 min, 70% B; 10 min, 50% B; 12 min, 50% B; 13 min, 25% B; 16 min, 25% B; 18 min, 0% B; 23 min, 0% B; 24 min, 85% B; 30 min, 85% B. Autosampler temperature was 5°C. Data were analyzed using the MAVEN software (Build # 682), Compound Discoverer software (Thermofisher Scientific), and R software. To control for instrument variability, an internal control [13C5, 15N]-valine, was spiked in the extraction solvent.

**References for Supplementary Materials and Methods**

1. Peng H, Carrico D, Thai V, Blaskovich M, Bucher C, Pusateri EE, et al. Synthesis and evaluation of potent, highly-selective, 3-aryl-piperazinone inhibitors of protein geranylgeranyltransferase-I. Org Biomol Chem. 2006 May 7;4(9):1768–84.

2. Jung SM, Doxsey WG, Le J, Haley JA, Mazuecos L, Luciano AK, et al. In vivo isotope tracing reveals the versatility of glucose as a brown adipose tissue substrate. Cell Rep. 2021 Jul 27;36(4):109459.

**Supplementary Figure Legends**

**Figure S1**: (A) Pitavastatin has cytotoxic activity at 24hr that is prevented by caspase inhibitor QVD-OPH. (B) PIT retains cytotoxic activity in VEN-resistant MOLM13-R cells. (C) Simvastatin is less potent in AML cell lines, with effective concentratons of 3 or 10 µM. Data are expressed as mean +/- SEM *p < 0.05, ** p < 0.01, ***p < 0.001, ****p < 0.0001 two-way ANOVA, n = 3.

**Figure S2**: (A, B) Additional viability assays on primary samples treated with pitavastatin alone (A) or in combination with venetoclax (B). (C) PIT increases cytotoxicity of VEN-AZA (5Aza) combo in primary AML samples. Data are expressed as mean +/- SEM. Genetic lesions are shown in parentheses.

**Figure S3**: PIT cytotoxicity and VEN sensitization is preserved in isogenic MOLM14 cell lines with *TP53* mutation. Graphs showing percentage of Annexin V and PI negative cells after treatment with different concentrations of pitavastatin in combination with venetoclax. Data are expressed as mean +/- SEM *p < 0.05, ** p < 0.01, ***p < 0.001, ****p < 0.0001. Two-way ANOVA with Tukey post hoc test, n ≥ 3.

**Figure S4**: PUMA upregulation in additional statin-sensitive and resistant cell lines. (A-B) Western blots were used to measure PUMA expression in pitavastatin sensitive (OCI-AML3, HNT-34) and insensitive (OCI-M1) AML cell lines. The PL-21 cell line is designated “weakly sensitive” because PIT alone has no cytotoxic effect but slightly enhances venetoclax cytotoxicity. Relative protein quantification performed by densitometric analysis using ImageJ64 software. Data are expressed as mean +/- SEM *p < 0.05, ** p < 0.01, ***p < 0.001, ****p < 0.0001. Two-way ANOVA, n ≥ 3. Similar experiments were performed to measure PUMA expression in an additional pitavastatin sensitive cell line, MV4-11 (C) and in MOLM13 where pitavastatin alone has no cytotoxic effect but sensitizes to venetoclax (D). Data are expressed as mean +/- SEM *p < 0.05, ** p < 0.01. One way ANOVA, n ≥ 3.

**Figure S5**: (A) Western blots were used to measure PUMA expression in control and *TP53* mutant MOLM14 isogenic cell lines. Data are expressed as mean +/- SEM *p < 0.05, ** p < 0.01, two-way ANOVA, n = 3. (B) Western blot validation of reduced or absent PUMA expression in OCI-AML3, MOLM13 and MOLM14 cells in which the *BBC3* gene was disrupted by CRISPR. Cells were treated with vehicle or pitavastatin (1 μM) for 16hr.

**Figure S6**: Pitavastatin treatment reduces or prevents the increase of MCL-1 protein expression in combination with venetoclax treatment. (A) Representative western blot of MCL-1 and PUMA protein expression in OCI-AML3, MOLM13, and MOLM14 cells following 24-hour treatments with Veh (0.1% DMSO), PIT, VEN and their combination. An antibody to vinculin was used as a loading control. (B) Quantification of relative MCL-1 and PUMA protein expression following treatment. Data are expressed as mean +/- SEM *p < 0.05, ** p < 0.01, ***p < 0.001, ****p < 0.0001 two-way ANOVA, n = 3. (C, D) PUMA and MCL1 protein analysis in MOLM13 parental and venetoclax resistant cell lines (C) and in three primary p53 mutant patient samples (D).

**Figure S7:** Data supporting pitavastatin mechanism is dependent on GGPP depletion. (A) GGPP rescue of cytotoxicity in sensitive lines (OCI-AML3, HNT-34). Data are expressed as mean +/- SEM, ** $p < 0.01$, ****$p < 0.0001$ two-way ANOVA (n = 2-4). (B) PUMA expression and relative quantification in OCI-AML3 after treatment with PIT, GGTI-298 or GGPS inhibitor (THZ145). Data are expressed as mean +/- SEM *$p < 0.05$, ** $p < 0.01$, one-way ANOVA, $n \geq 3$. (C) Graphs showing percentage of viable OCI-AML3 after 48h of treatment with pitavastatin 1 μM, GGTI-298, GGTI-2133 or THZ145 in combination with venetoclax. Data are expressed as mean +/- SEM **** $p < 0.0001$, two-way ANOVA, n =3.

**Figure S8**: GGTIs inhibit prenylation. (A) F-actin polymeration measured by phalloidin staining is inhibited by both GGTI-298 and GGTI-2133. Latrunculin was used as a positive control to completely disrupt actin polymerization. (B, C) All three GGTIs cause RAP1A unprenylation; GGTI-2417 increases PUMA to a similar or greater extent as GGTI-298. (D) percentage of viable OCI-AML3 treated with pitavastatin, GGTI-298 or GGTI-2417 in combination with venetoclax. Data are expressed as mean +/- SEM *$p < 0.05$, ** $p < 0.01$, ****$p < 0.0001$. Two-way ANOVA, n = 4.

**Figure S9**: (A) Venn diagram and intersection plots for HNT-34 cells, similar to Figure 3A for OCI-AML3 cells. (B) PCA plots for transcriptome experiment #1. (C) Intersection plots of sample distance for OCI-AML3 (left) and HNT-34 (right). (D) Additional evidence for MYC/E2F downregulation using Ingenuity Pathway Analysis.

**Figure S10**: Fold change in mRNA expression of genes of interest described in the Results section. (A) OCI-AML3 cells. (B) HNT-34 cells. (C) OCI-AML3 cells from experiment #2.

**Figure S11:** (A) PCA analysis of OCI-AML3 cells analyzed in transcriptome experiment #2. (B) Venn diagram and intersection plot for MOLM13 cells in experiment #2. Consistent with minimal pitavastatin sensitivity, these cells had few DEGs. (C) GSEA analysis of data from MOLM13 cells identified cholesterol and lipid pathways as predominant gene sets.

**Figure S12**: MYC downregulation in OCI-AML3 cells is recapitulated by GGTI-298 and GGTI-2417, weakly by GGTI-2133. (A) Representative western blot. (B) Relative protein quantifications. Data are expressed as mean +/- SEM *p < 0.05, ** p < 0.01, *** p < 0.001 one-way ANOVA, n = 3.

**Figure S13**: CRISPR KO of the *TSC22D3* gene encoding GILZ does not alter PIT cytotoxicity or combo effect with VEN. (A) GILZ upregulation and MYC downregulation are rescued by GGPP. Top: representative Western blot. MM1S cells are a dexamethasone (Dex)-sensitive MM cell line used as a positive control for GILZ induction. Bottom: graphs depict quantitation of 3 experiments. Data are expressed as mean +/- SEM *p < 0.05, ** p < 0.01, **** p < 0.0001 two-way ANOVA. (B) Western blot showing loss of GILZ expression in knockout clones. (C) Viability assay in parental, empty vector (EV) control, and three knockout clones.

**Figure S14**: Mitochondrial mass was measured in AML cell lines treated with vehicle or pitavastatin for 16hr. (A) TOM20 measured by intracellular staining and flow cytometry. (B) Mitochondrial DNA copy number relative to genomic DNA was determined by qPCR. (C)

Cardiolipin measured by intracellular staining and flow cytometry. Data are expressed as mean +/- SEM *p < 0.05, ***p < 0.001 two-way ANOVA.

**Figure S15:** (A-B) Graphs showing reduction of maximal OCR and spare respiratory capacity by the combination of PIT + VEN in OCI-AML3 and MOLM13. Data are expressed as mean +/- SEM *p < 0.05, ** p < 0.01, ***p < 0.001, **** p < 0.0001 two-way ANOVA. (C, D) Flow cytometry was used to measure the percentage of cells that release Cytochrome c (Cyt-c) after 16hr of treatment. (C) In OCI-AML3 and MOLM13 cells, venetoclax alone causes Cyt-c release that is enhanced by PIT in a GGPP-dependent manner. (D) PIT + VEN causes Cyt-c release in control and *TP53* mutant MOLM14 cells. Data are expressed as mean of cytochrome-C-high cells, +/- SEM, *p < 0.05, ** p < 0.01, ****p < 0.0001 two-way ANOVA, n = 3.

**Figure S16:** Pitavastatin causes loss of mitochondrial membrane potential (MMP) in a significant fraction of OCI-AML3 (A) and MOLM14 cells including *TP53*-mutant (B). MMP loss was measured by flow cytometry using TMRE staining and expressed as percent MMP-low. Data are expressed as mean +/- SEM. *p < 0.05, ** p < 0.01, by two-way ANOVA, n = 5.

**Table S1:** Synergy scores for cell lines treated with pitavastatin/venetoclax combinations.

**Table S2:** Synergy scores for MOLM14 and derivative cell lines treated with pitavastatin/venetoclax combinations.

**Supplementary Data Files:**

**Excel File #1: DEGs lists and volcano plots for transcriptome experiment #1.**

**Excel File #2: DEGs lists and volcano plots for transcriptome experiment #2.**

DEG lists were generated using an adjusted p value threshold of 0.05 and a $\log_2$ fold-change (LFC) threshold of +/- 0.1. Volcano plots display DEGs with a more stringent LFC threshold for ease of visualization of named genes.

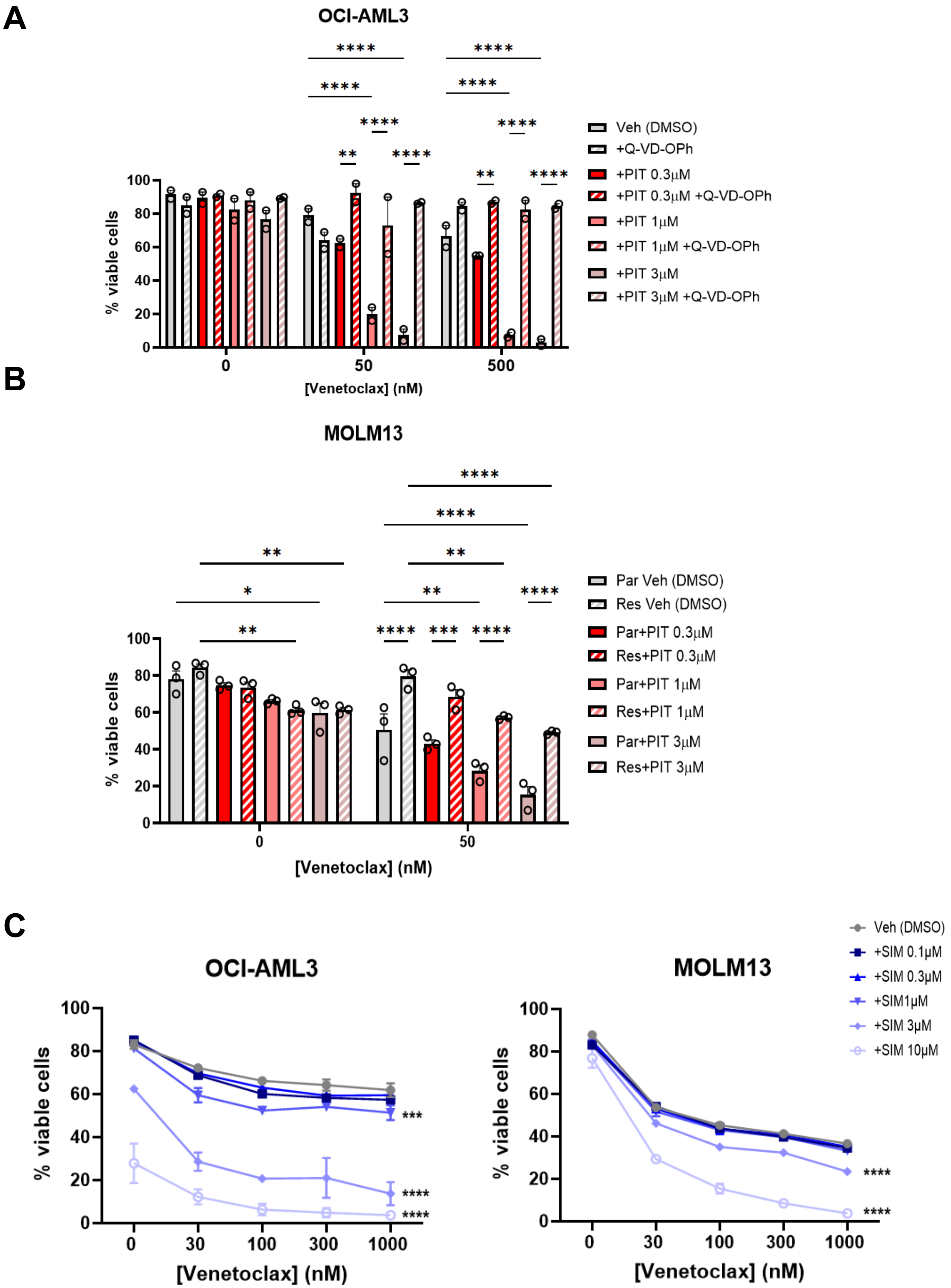
A
OCI-AML3
% viable cells
[Venetoclax] (nM)
Veh (DMSO)
+Q-VD-OPh
+PIT 0.3µM
+PIT 0.3µM +Q-VD-OPh
+PIT 1µM
+PIT 1µM +Q-VD-OPh
+PIT 3µM
+PIT 3µM +Q-VD-OPh
B
MOLM13
Par Veh (DMSO)
Res Veh (DMSO)
Par+PIT 0.3µM
Res+PIT 0.3µM
Par+PIT 1µM
Res+PIT 1µM
Par+PIT 3µM
Res+PIT 3µM
C
Veh (DMSO)
+SIM 0.1µM
+SIM 0.3µM
+SIM1µM
+SIM 3µM
+SIM 10µM

Fig. S1

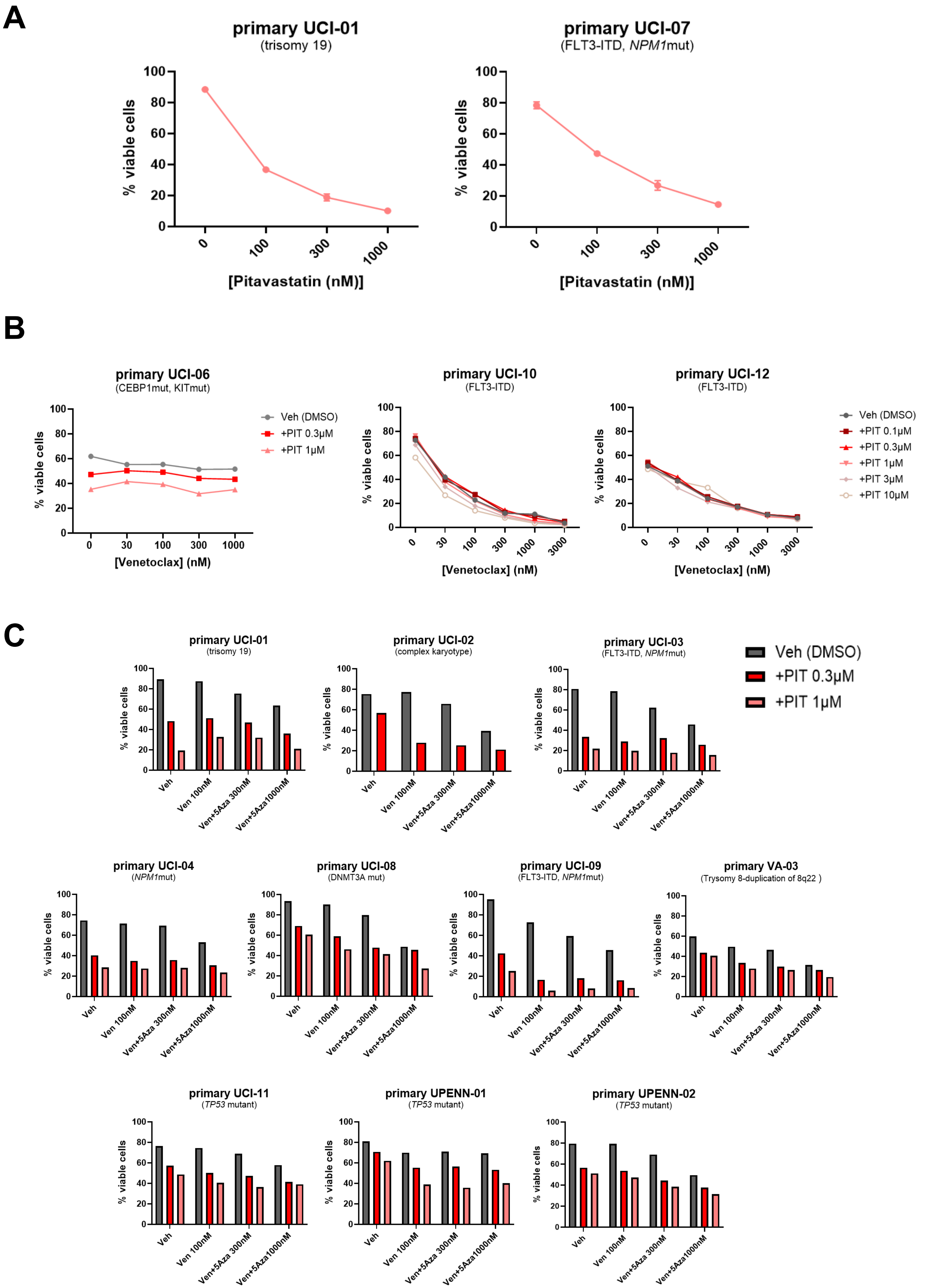
A
primary UCI-01
(trisomy 19)
primary UCI-07
(FLT3-ITD, NPM1mut)
% viable cells
[Pitavastatin (nM)]
B
primary UCI-06
(CEBP1mut, KITmut)
primary UCI-10
(FLT3-ITD)
primary UCI-12
(FLT3-ITD)
Veh (DMSO)
+PIT 0.1μM
+PIT 0.3μM
+PIT 1μM
+PIT 3μM
+PIT 10μM
[Venetoclax] (nM)
C
primary UCI-01
(trisomy 19)
primary UCI-02
(complex karyotype)
primary UCI-03
(FLT3-ITD, NPM1mut)
primary UCI-04
(NPM1mut)
primary UCI-08
(DNMT3A mut)
primary UCI-09
(FLT3-ITD, NPM1mut)
primary VA-03
(Trysomy 8-duplication of 8q22 )
primary UCI-11
(TP53 mutant)
primary UPENN-01
(TP53 mutant)
primary UPENN-02
(TP53 mutant)
Veh
Ven 100nM
Ven+5Aza 300nM
Ven+5Aza1000nM

Fig. S2

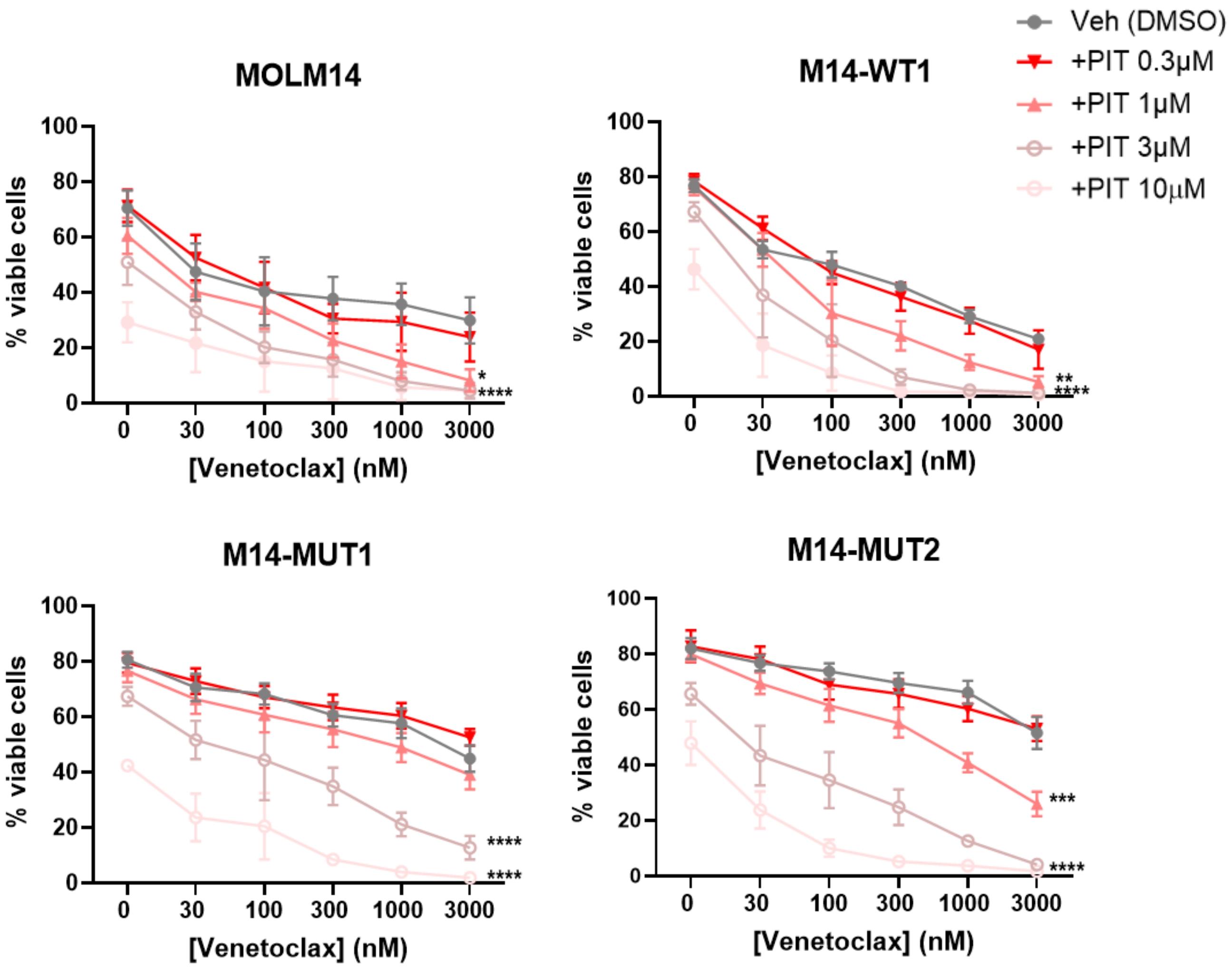
Veh (DMSO)
+PIT 0.3μM
+PIT 1μM
+PIT 3μM
+PIT 10μM
MOLM14
% viable cells
[Venetoclax] (nM)
*
****
M14-WT1
**
****
M14-MUT1
****
****
M14-MUT2
***
****

Fig S3

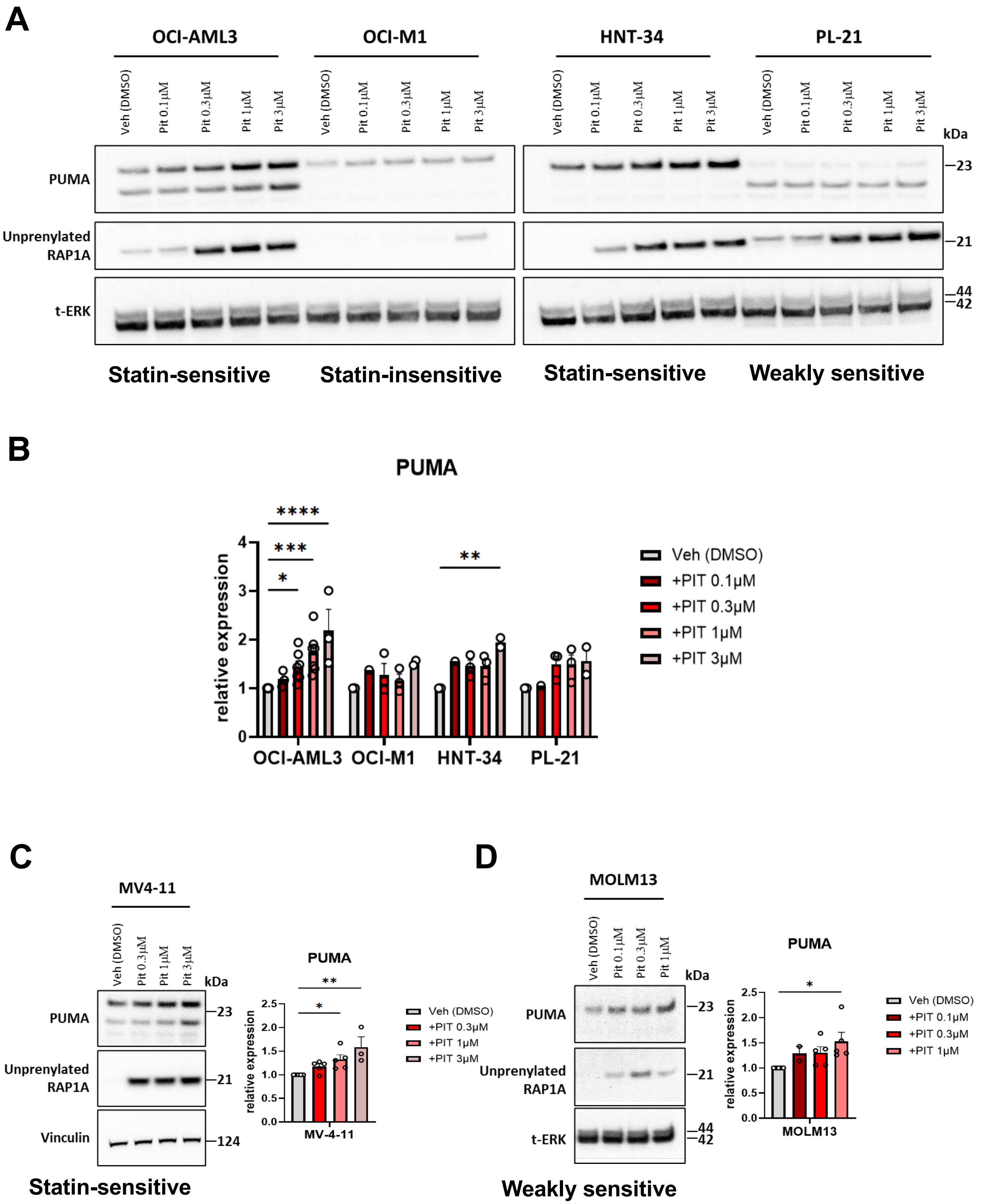

A
OCI-AML3
OCI-M1
HNT-34
PL-21
Veh (DMSO)
Pit 0.1µM
Pit 0.3µM
Pit 1µM
Pit 3µM
kDa
PUMA
Unprenylated RAP1A
t-ERK
23
21
44
42
Statin-sensitive
Statin-insensitive
Statin-sensitive
Weakly sensitive
B
PUMA
relative expression
OCI-AML3
OCI-M1
HNT-34
PL-21
Veh (DMSO)
+PIT 0.1µM
+PIT 0.3µM
+PIT 1µM
+PIT 3µM
C
MV4-11
PUMA
Unprenylated RAP1A
Vinculin
124
MV-4-11
Statin-sensitive
D
MOLM13
Weakly sensitive


Fig. S4

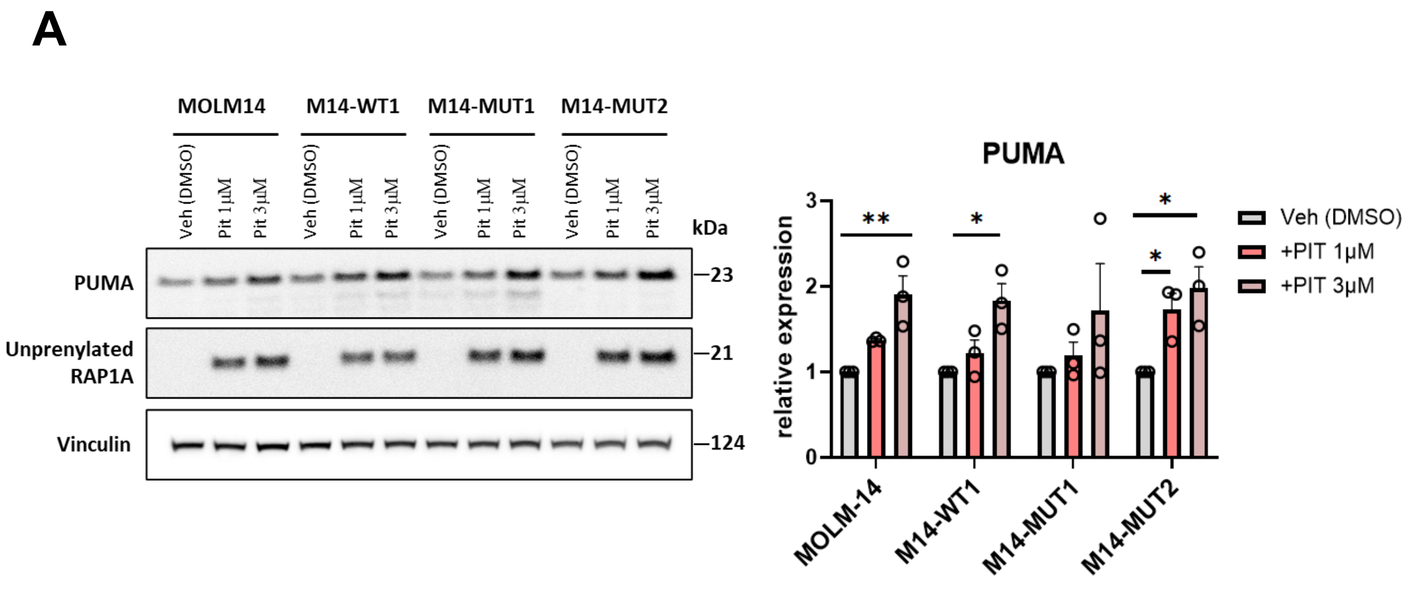
A
MOLM14
M14-WT1
M14-MUT1
M14-MUT2
Veh (DMSO)
Pit 1µM
Pit 3µM
kDa
PUMA
Unprenylated RAP1A
Vinculin
23
21
124
PUMA
relative expression
Veh (DMSO)
+PIT 1µM
+PIT 3µM
MOLM-14
M14-WT1
M14-MUT1
M14-MUT2

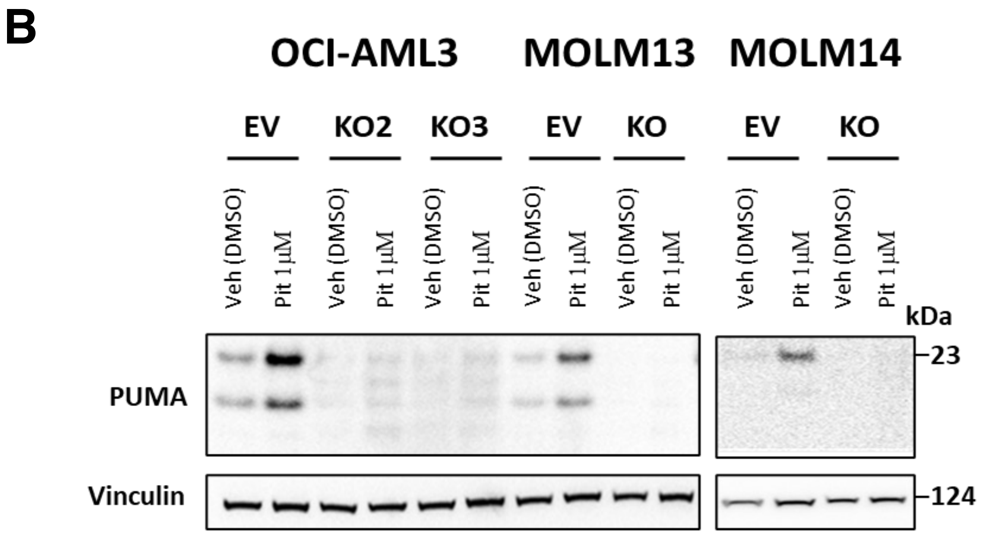
B
OCI-AML3
MOLM13
MOLM14
EV
KO2
KO3
KO
Veh (DMSO)
Pit 1µM
kDa
PUMA
Vinculin
23
124

Fig. S5

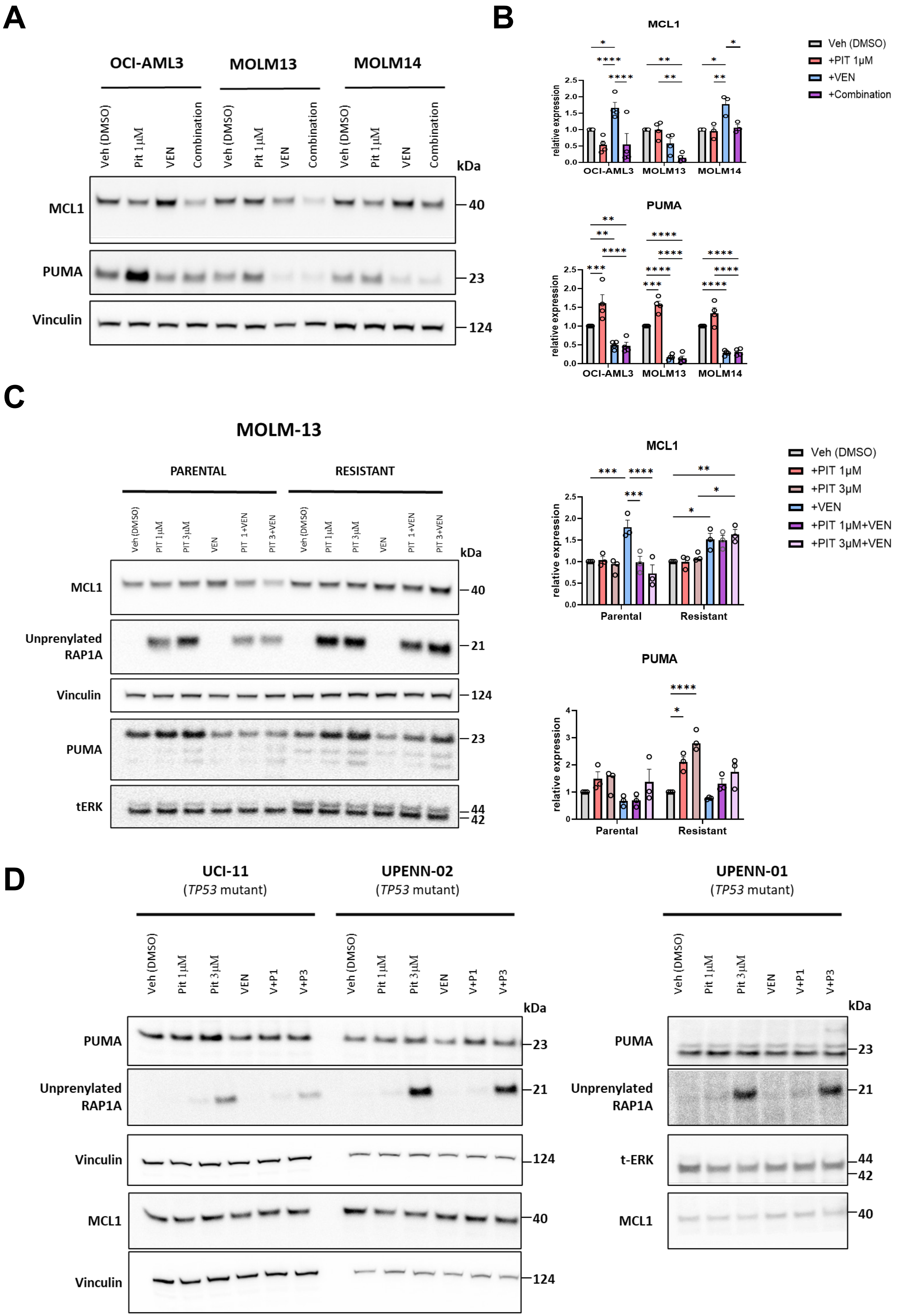
A
OCI-AML3
MOLM13
MOLM14
Veh (DMSO)
Pit 1 μM
VEN
Combination
kDa
MCL1
PUMA
Vinculin
40
23
124
B
MCL1
PUMA
relative expression
OCI-AML3
MOLM13
MOLM14
Veh (DMSO)
+PIT 1μM
+VEN
+Combination
C
MOLM-13
PARENTAL
RESISTANT
Veh (DMSO)
PIT 1μM
PIT 3μM
VEN
PIT 1+VEN
PIT 3+VEN
kDa
MCL1
Unprenylated RAP1A
Vinculin
PUMA
tERK
40
21
124
23
44
42
MCL1
PUMA
relative expression
Parental
Resistant
Veh (DMSO)
+PIT 1μM
+PIT 3μM
+VEN
+PIT 1μM+VEN
+PIT 3μM+VEN
D
UCI-11
(TP53 mutant)
UPENN-02
(TP53 mutant)
UPENN-01
(TP53 mutant)
Veh (DMSO)
Pit 1 μM
Pit 3 μM
VEN
V+P1
V+P3
kDa
PUMA
Unprenylated RAP1A
Vinculin
MCL1
t-ERK
23
21
124
40
44
42

Fig. S6

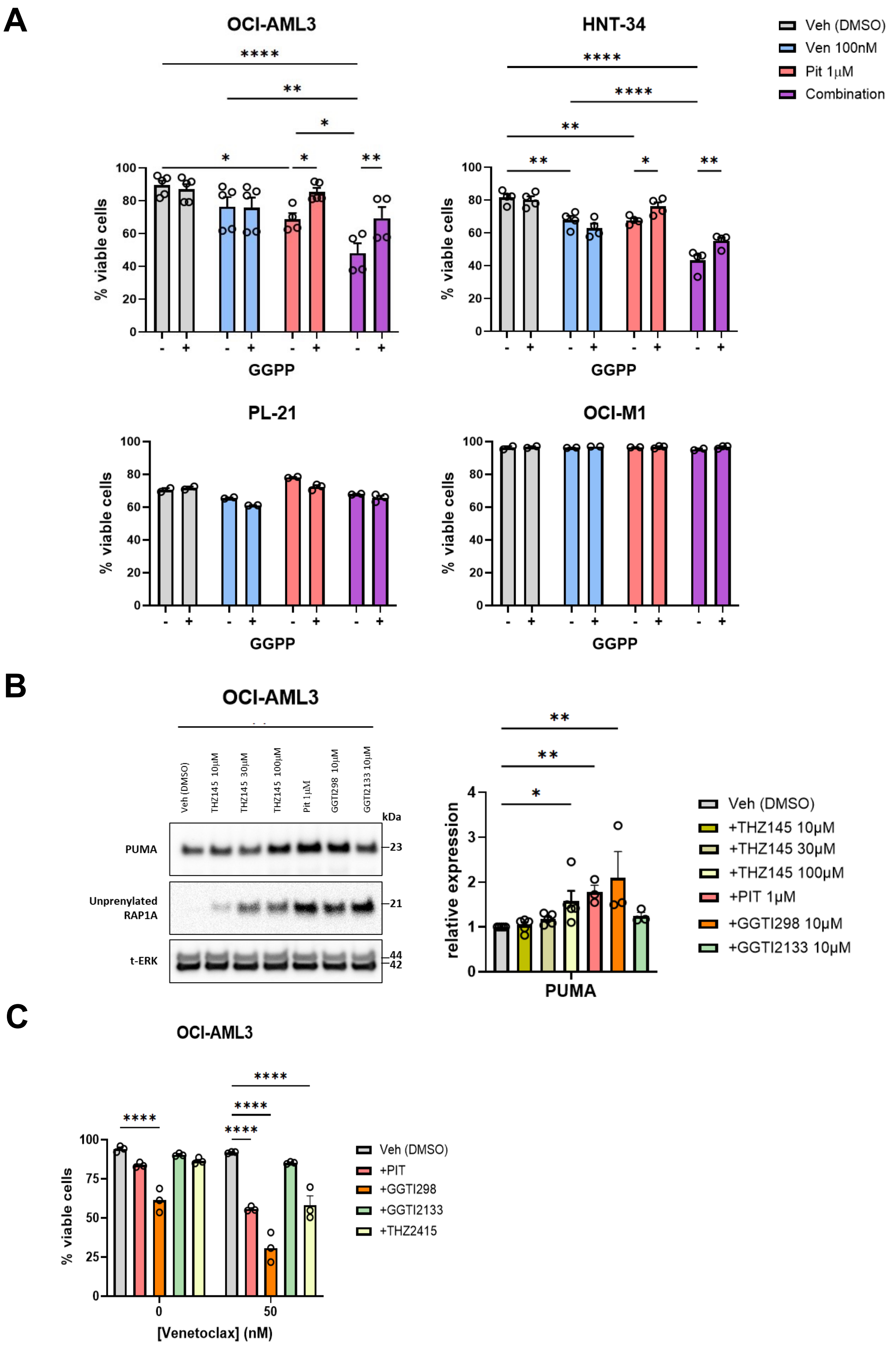
A
OCI-AML3
HNT-34
Veh (DMSO)
Ven 100nM
Pit 1µM
Combination
% viable cells
GGPP
PL-21
OCI-M1
B
OCI-AML3
Veh (DMSO)
THZ145 10µM
THZ145 30µM
THZ145 100µM
Pit 1µM
GGTI298 10µM
GGTI2133 10µM
kDa
PUMA
Unprenylated RAP1A
t-ERK
23
21
44
42
relative expression
PUMA
+THZ145 10µM
+THZ145 30µM
+THZ145 100µM
+PIT 1µM
+GGTI298 10µM
+GGTI2133 10µM
C
OCI-AML3
% viable cells
[Venetoclax] (nM)
+PIT
+GGTI298
+GGTI2133
+THZ2415

Fig. S7

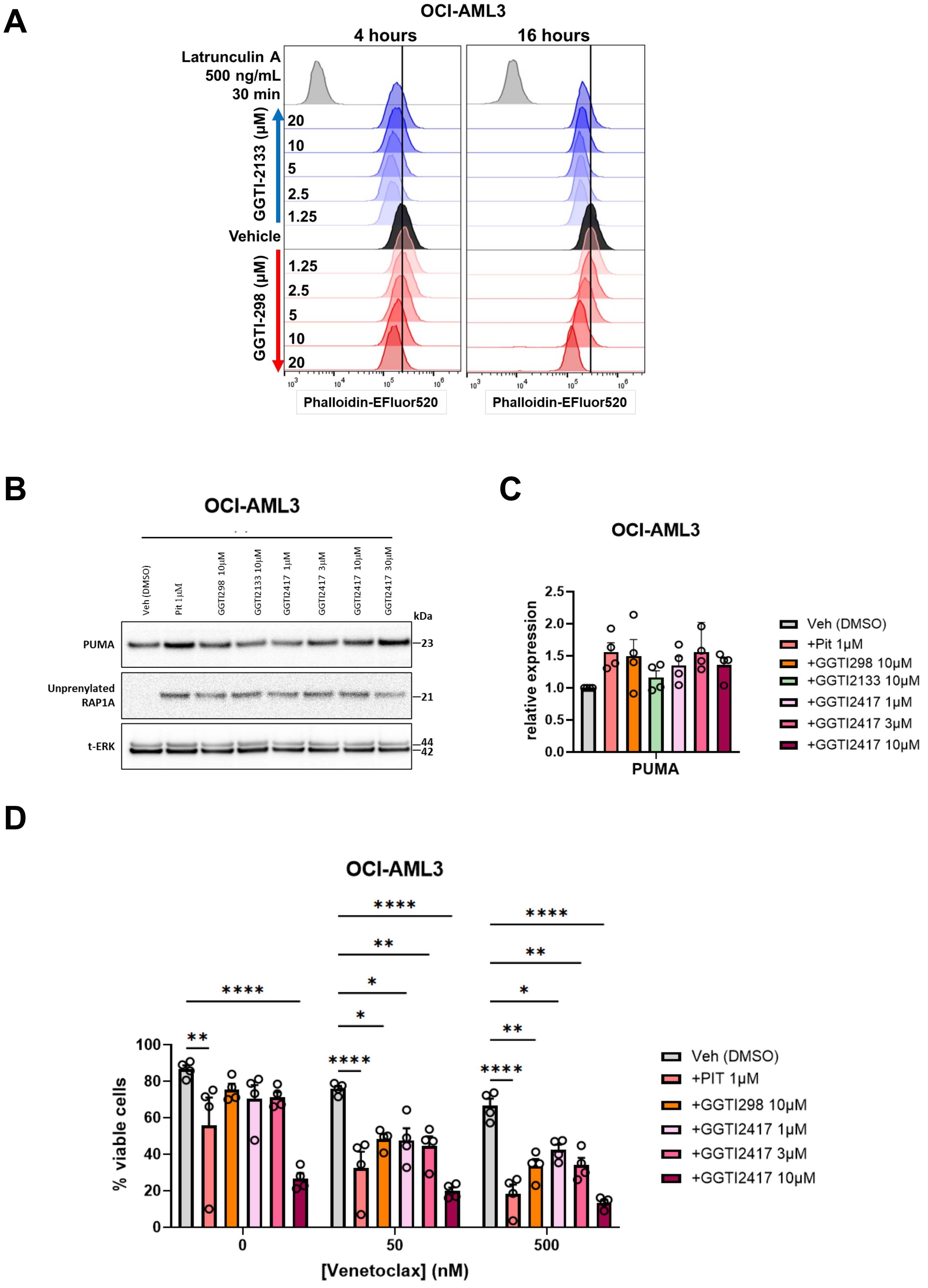
A
OCI-AML3
4 hours
16 hours
Latrunculin A
500 ng/mL
30 min
GGTI-2133 (μM)
20
10
5
2.5
1.25
Vehicle
GGTI-298 (μM)
1.25
2.5
5
10
20
Phalloidin-EFluor520
Phalloidin-EFluor520
B
OCI-AML3
Veh (DMSO)
Pit 1μM
GGTI298 10μM
GGTI2133 10μM
GGTI2417 1μM
GGTI2417 3μM
GGTI2417 10μM
GGTI2417 30μM
kDa
PUMA
23
Unprenylated
RAP1A
21
t-ERK
44
42
C
OCI-AML3
relative expression
PUMA
Veh (DMSO)
+Pit 1μM
+GGTI298 10μM
+GGTI2133 10μM
+GGTI2417 1μM
+GGTI2417 3μM
+GGTI2417 10μM
D
OCI-AML3
% viable cells
[Venetoclax] (nM)
Veh (DMSO)
+PIT 1μM
+GGTI298 10μM
+GGTI2417 1μM
+GGTI2417 3μM
+GGTI2417 10μM

Fig. S8

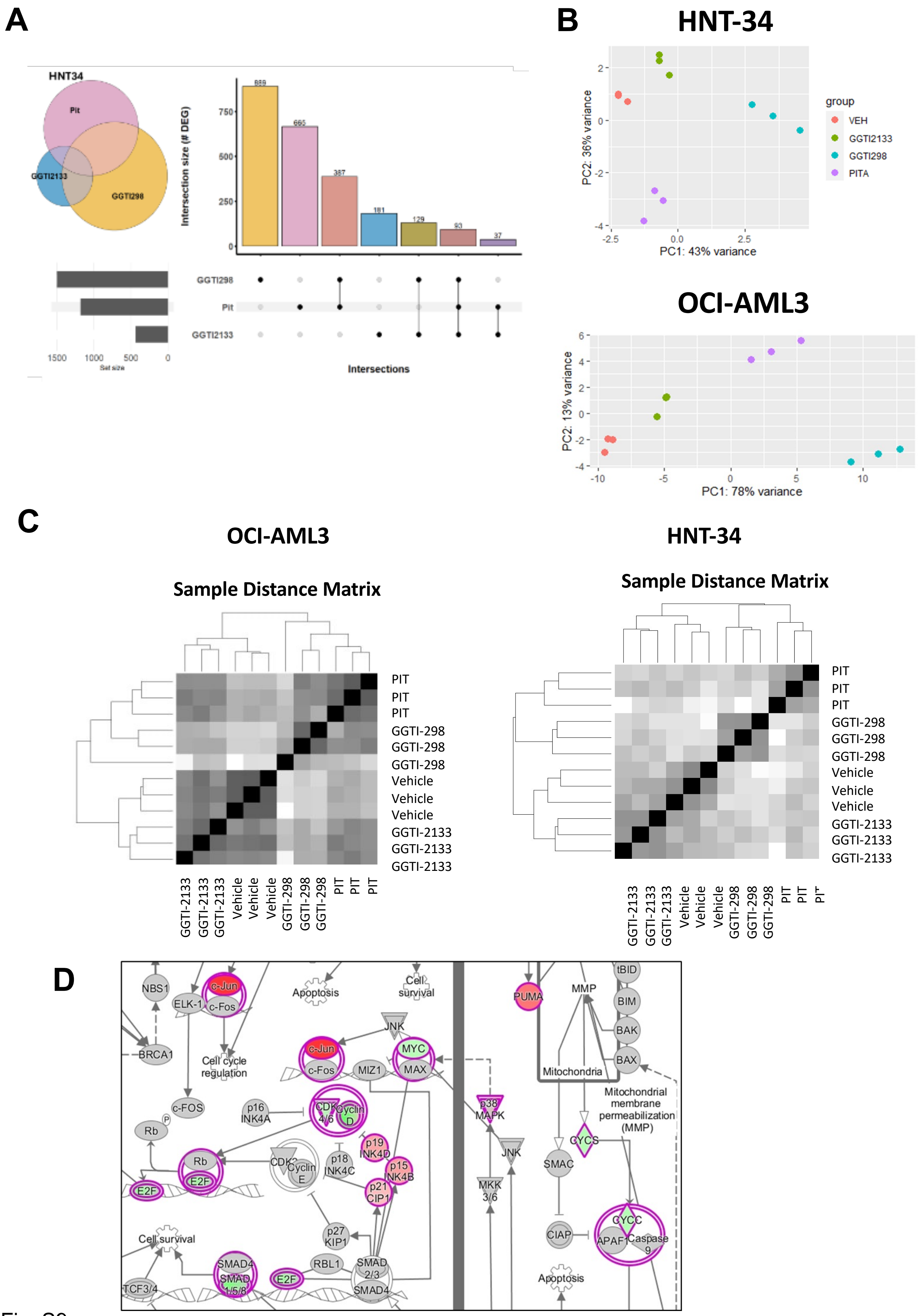
A
HNT34
Pit
GGTI2133
GGTI298
Intersection size (# DEG)
889
665
387
181
129
93
37
GGTI298
Pit
GGTI2133
1500 1000 500 0
Set size
Intersections
B
HNT-34
PC2: 36% variance
PC1: 43% variance
group
VEH
GGTI2133
GGTI298
PITA
OCI-AML3
PC2: 13% variance
PC1: 78% variance
C
OCI-AML3
Sample Distance Matrix
PIT
GGTI-298
Vehicle
GGTI-2133
HNT-34
Sample Distance Matrix
D
NBS1
c-Jun
c-Fos
ELK-1
BRCA1
Apoptosis
Cell survival
Cell cycle regulation
JNK
MYC
MAX
MIZ1
c-FOS
p16 INK4A
CDK 4/6
Cyclin D
Rb
E2F
p19 INK4D
p18 INK4C
p15 INK4B
p21 CIP1
p27 KIP1
Cyclin E
SMAD4
SMAD 1/5/8
SMAD 2/3
RBL1
TCF3/4
PUMA
MMP
tBID
BIM
BAK
BAX
Mitochondria
Mitochondrial membrane permeabilization (MMP)
p38 MAPK
MKK 3/6
CYCS
SMAC
CIAP
CYCC
APAF1
Caspase 9

Fig. S9

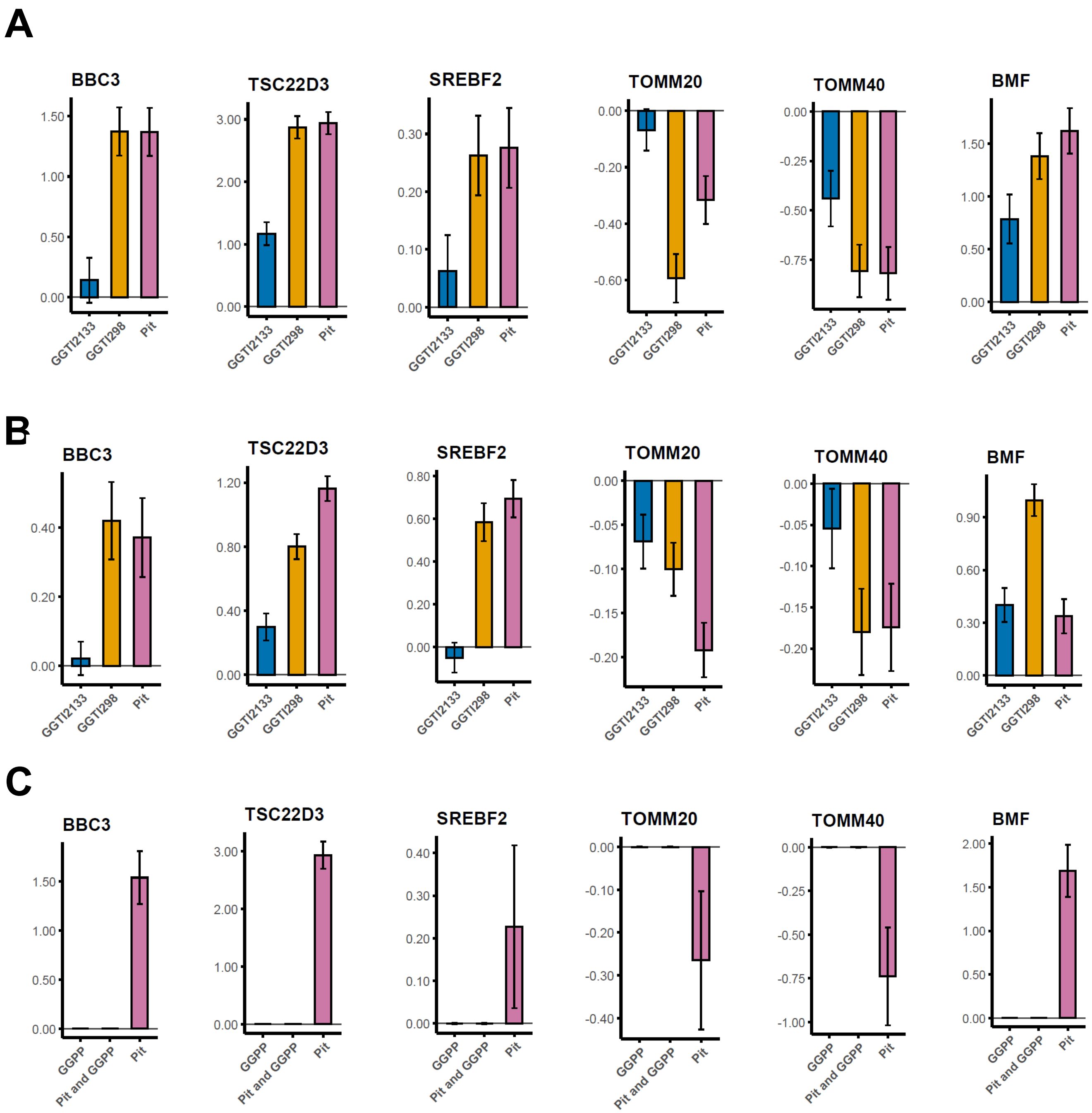
A
BBC3
TSC22D3
SREBF2
TOMM20
TOMM40
BMF
GGTI2133
GGTI298
Pit
B
C
GGPP
Pit and GGPP

Fig. S10

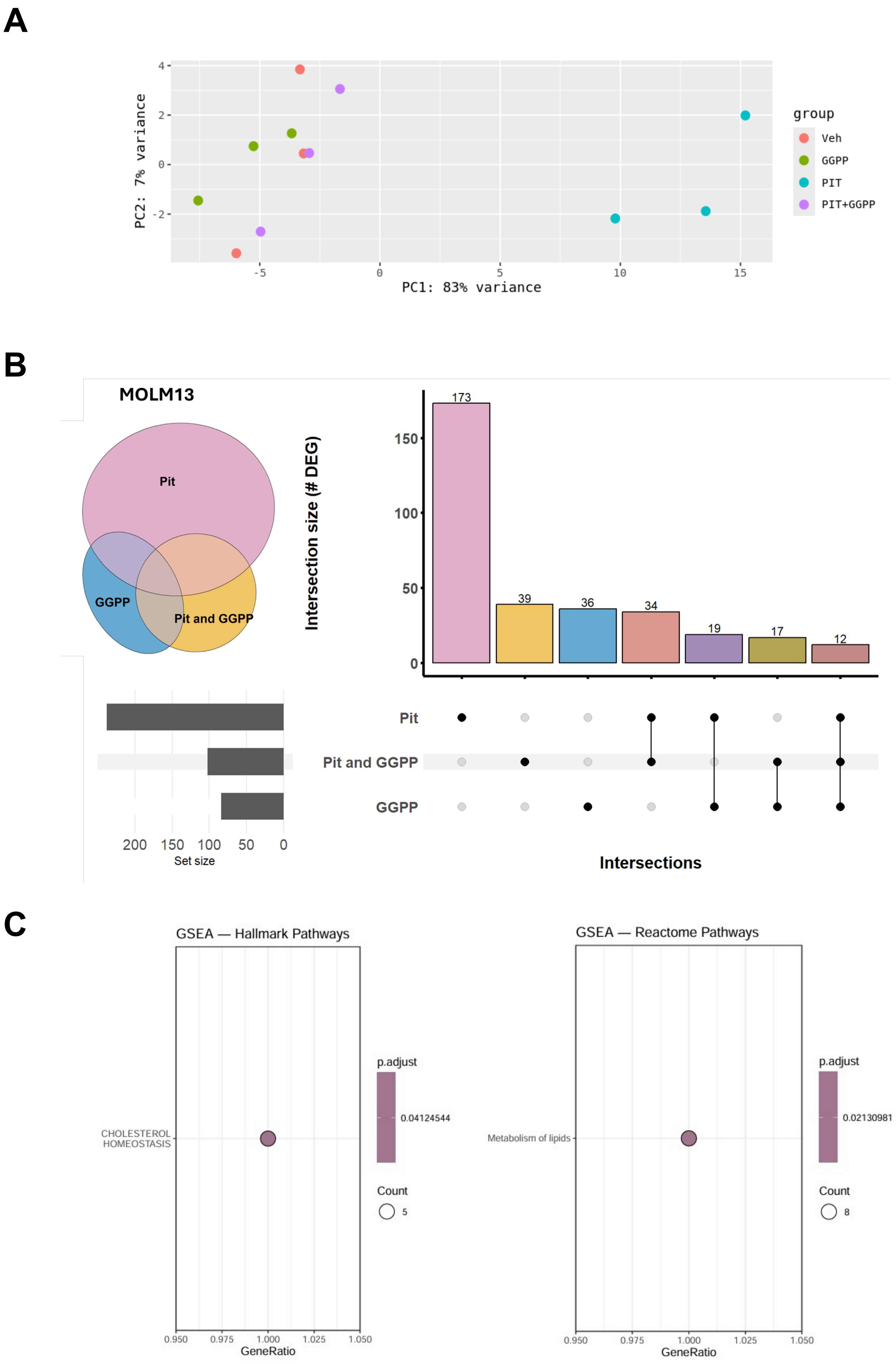
A
PC2: 7% variance
PC1: 83% variance
group
Veh
GGPP
PIT
PIT+GGPP
B
MOLM13
Pit
GGPP
Pit and GGPP
Intersection size (# DEG)
173
39
36
34
19
17
12
Set size
Intersections
C
GSEA — Hallmark Pathways
CHOLESTEROL HOMEOSTASIS
p.adjust
0.04124544
Count
5
GeneRatio
GSEA — Reactome Pathways
Metabolism of lipids
p.adjust
0.02130981
Count
8
GeneRatio

Fig. S11

A

B

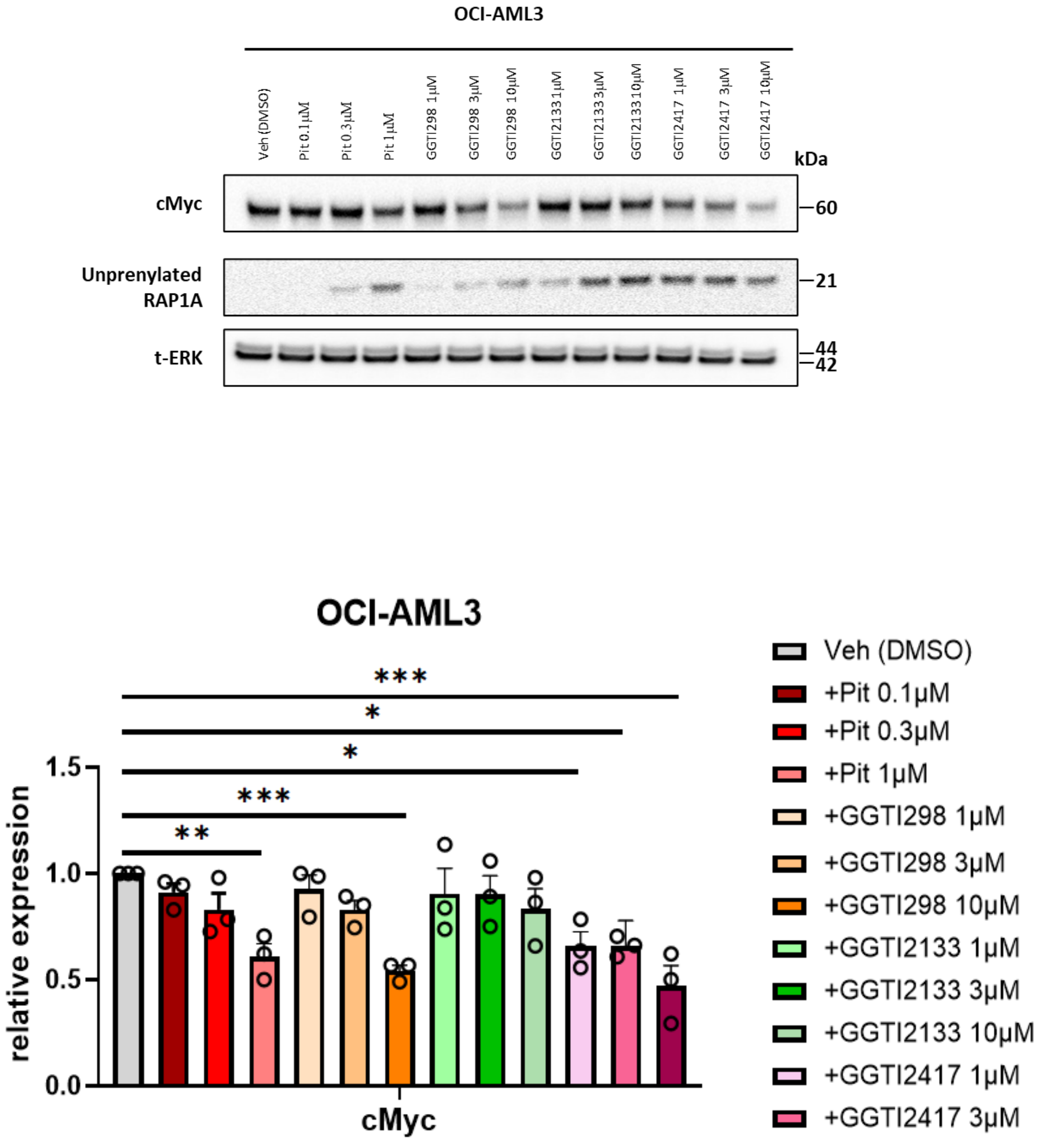

OCI-AML3
Veh (DMSO)
Pit 0.1µM
Pit 0.3µM
Pit 1µM
GGTI298 1µM
GGTI298 3µM
GGTI298 10µM
GGTI2133 1µM
GGTI2133 3µM
GGTI2133 10µM
GGTI2417 1µM
GGTI2417 3µM
GGTI2417 10µM
kDa
cMyc
60
Unprenylated RAP1A
21
t-ERK
44
42
OCI-AML3
***
*
*
***
**
1.5
1.0
0.5
0.0
relative expression
cMyc
Veh (DMSO)
+Pit 0.1µM
+Pit 0.3µM
+Pit 1µM
+GGTI298 1µM
+GGTI298 3µM
+GGTI298 10µM
+GGTI2133 1µM
+GGTI2133 3µM
+GGTI2133 10µM
+GGTI2417 1µM
+GGTI2417 3µM
+GGTI2417 10µM


Fig. S12

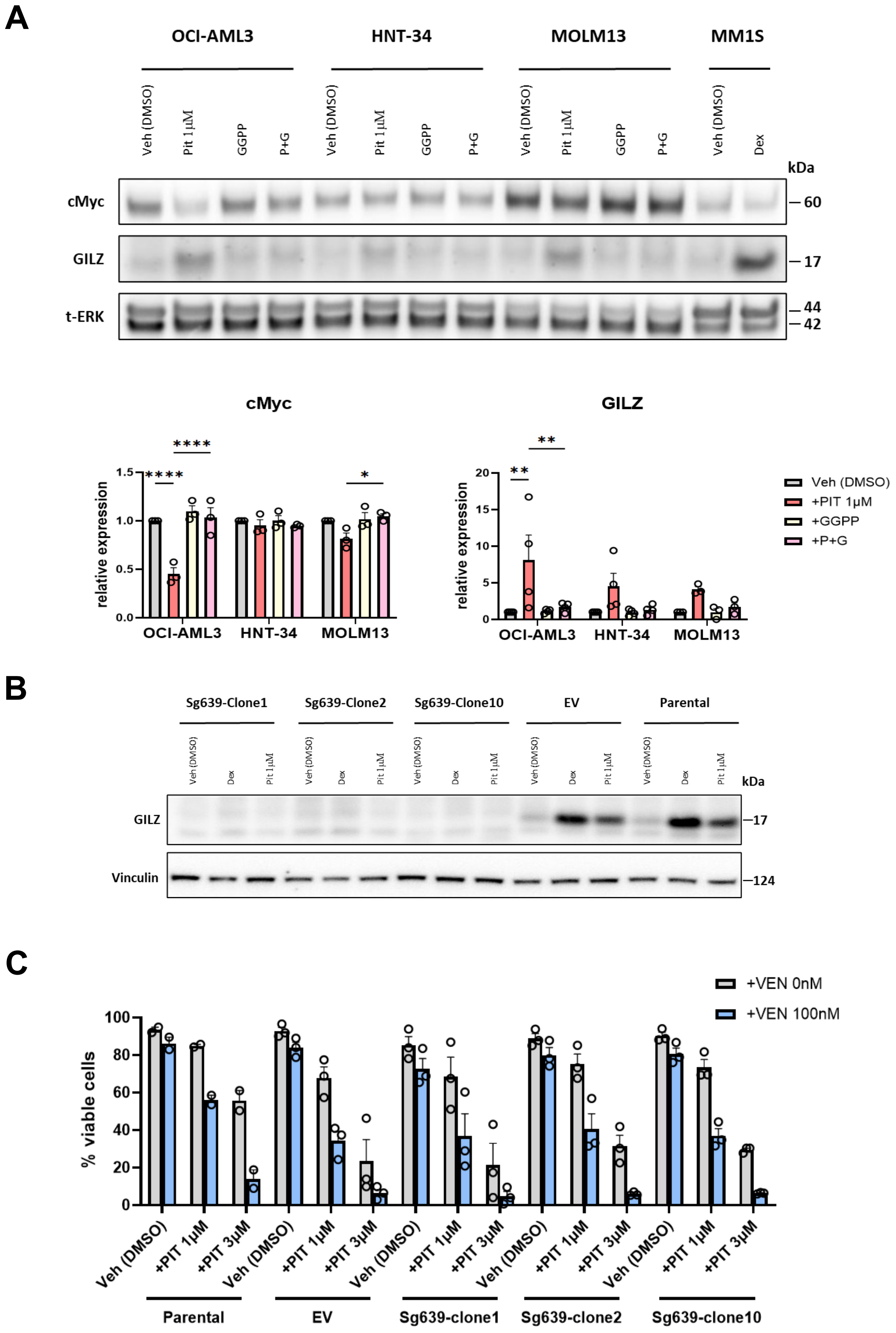

A
OCI-AML3
HNT-34
MOLM13
MM1S
Veh (DMSO)
Pit 1μM
GGPP
P+G
Dex
kDa
cMyc
GILZ
t-ERK
60
17
44
42
cMyc
GILZ
relative expression
Veh (DMSO)
+PIT 1μM
+GGPP
+P+G
B
Sg639-Clone1
Sg639-Clone2
Sg639-Clone10
EV
Parental
Vinculin
124
C
+VEN 0nM
+VEN 100nM
% viable cells
+PIT 1μM
+PIT 3μM
Sg639-clone1
Sg639-clone2
Sg639-clone10


Fig. S13

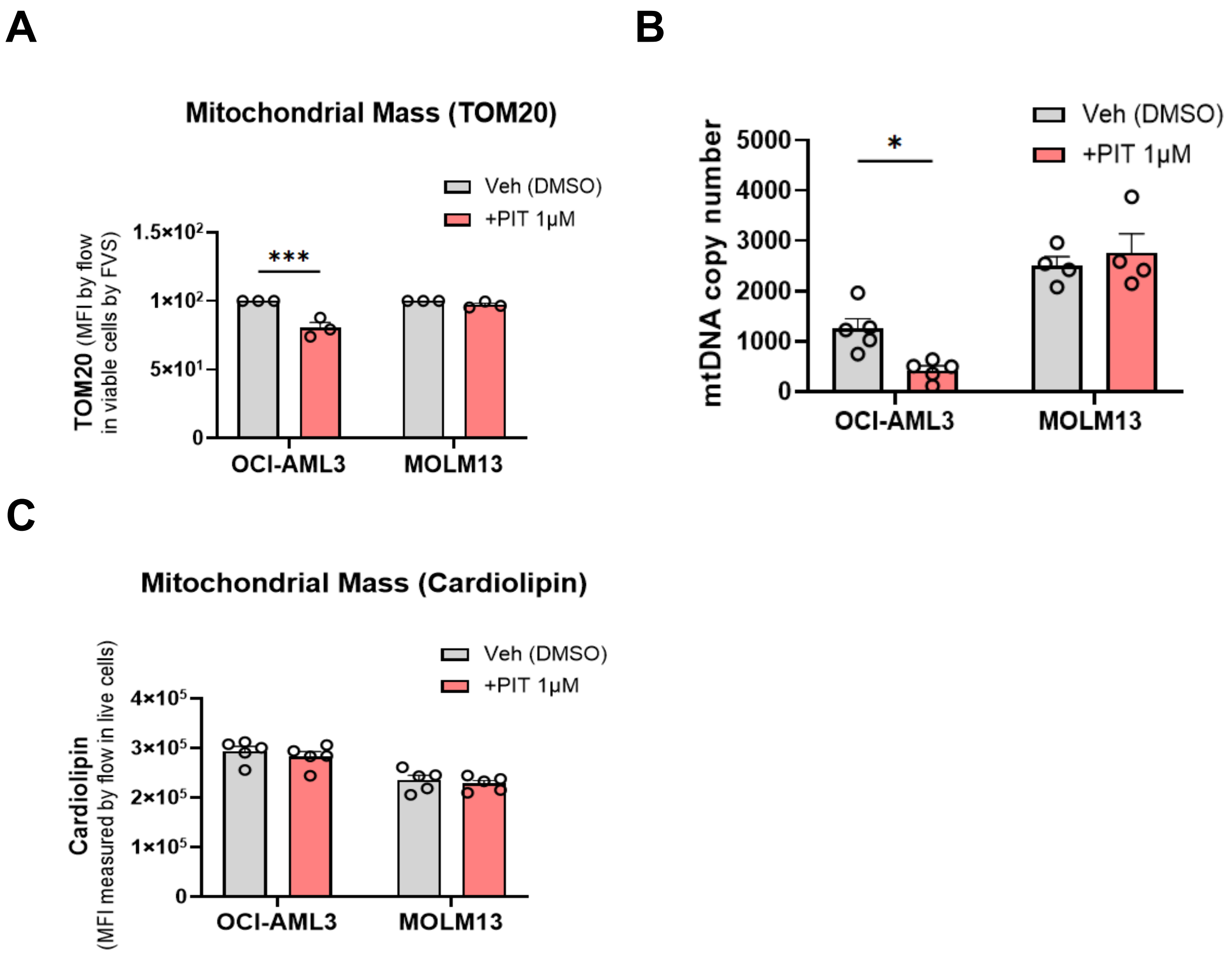
A
Mitochondrial Mass (TOM20)
Veh (DMSO)
+PIT 1μM
TOM20 (MFI by flow in viable cells by FVS)
1.5×10²
1×10²
5×10¹
0
***
OCI-AML3
MOLM13
B
Veh (DMSO)
+PIT 1μM
mtDNA copy number
5000
4000
3000
2000
1000
0
*
OCI-AML3
MOLM13
C
Mitochondrial Mass (Cardiolipin)
Veh (DMSO)
+PIT 1μM
Cardiolipin (MFI measured by flow in live cells)
4×10⁵
3×10⁵
2×10⁵
1×10⁵
0
OCI-AML3
MOLM13

Fig. S14

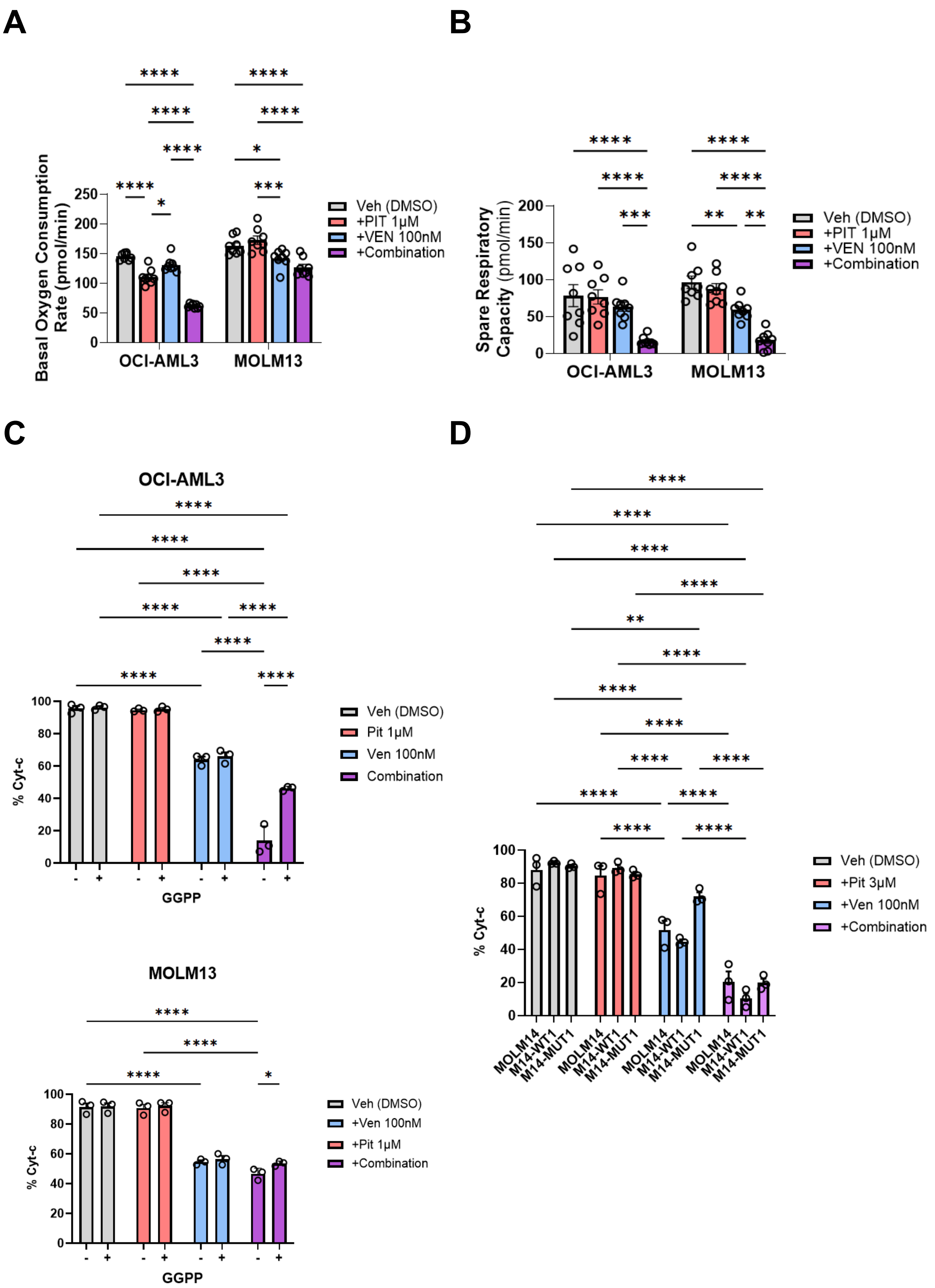
A
B
C
D
Basal Oxygen Consumption Rate (pmol/min)
OCI-AML3
MOLM13
Veh (DMSO)
+PIT 1μM
+VEN 100nM
+Combination
Spare Respiratory Capacity (pmol/min)
OCI-AML3
% Cyt-c
GGPP
Veh (DMSO)
Pit 1μM
Ven 100nM
Combination
MOLM13
+Ven 100nM
+Pit 1μM
+Pit 3μM
MOLM14
M14-WT1
M14-MUT1

Fig. S15

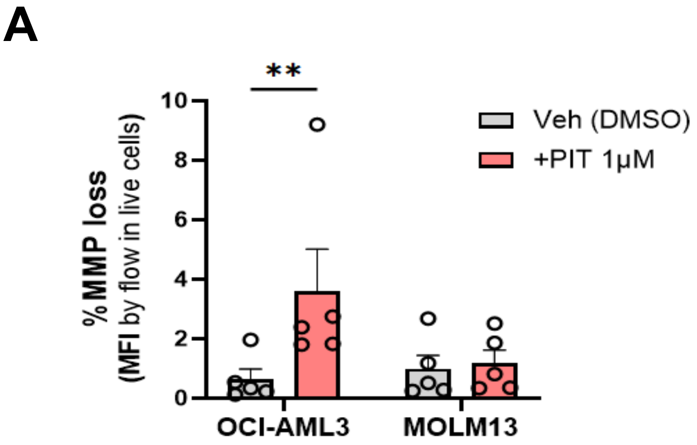
A
**
10
8
6
4
2
0
% MMP loss
(MFI by flow in live cells)
Veh (DMSO)
+PIT 1μM
OCI-AML3
MOLM13

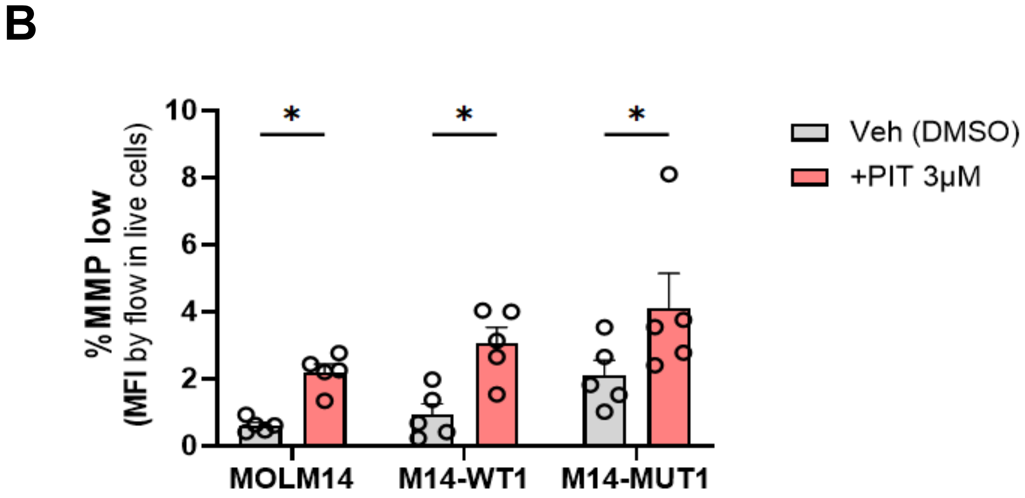
B
10
8
6
4
2
0
*
*
*
% MMP low
(MFI by flow in live cells)
Veh (DMSO)
+PIT 3μM
MOLM14
M14-WT1
M14-MUT1

Fig. S16

**Table S1**

Bliss and HSA (highest single agent) synergy score and the most synergistic area determined by SynergyFinder 3.0 web-application. Synergy scores are categorized as follows: a score below -10 indicates antagonism, a score between -10 and 10 suggests an additive effect, and a score above 10 signifies synergy.

**Bliss and HSA Synergy Score of different AML cell lines treated with Pitavastatin and Venetoclax**

| Cell line | Combination | Synergy Score (Bliss) | Most Synergist Area (Bliss) | Synergy Score (HSA) | Most Synergist Area (HSA) |
|---|---|---|---|---|---|
| **OCI-AML3** | VEN+PIT | 12.475 | 15.175 | 24.629 | 27.999 |
| **HNT-34** | VEN+PIT | 15.33 | 16.56 | 24.403 | 25.243 |
| **PL-21** | VEN+PIT | 4.857 | 8.547 | 10.528 | 14.234 |
| **OCI-M1** | VEN+PIT | -9.371 | -4.038 | -8.441 | -3.552 |
| **MOLM13** | VEN+PIT | 15.24 | 21.568 | 15.24 | 21.568 |
| **MV-4-11** | VEN+PIT | 9.783 | 18.53 | 12.291 | 23.221 |

**Table S2**
Bliss and HSA synergy score and the most synergistic area calculated by SynergyFinder 3.0 web-application. Data are expressed as mean +/- SEM (n ≥ 3).

**Bliss and HSA Synergy Score of *p53*-WT and mutant cell lines treated with Pitavastatin and Venetoclax**

| **Cell line** | **Combination** | **Synergy Score (Bliss)** | **Most Synergist Area (Bliss)** | **Synergy Score (HSA)** | **Most Synergist Area (HSA)** |
|---|---|---|---|---|---|
| **MOLM14** | VEN+PIT | 9.214 | 22.158 | 19.446 | 35.517 |
| **M14-WT1** | VEN+PIT | 20.909 | 31.402 | 26.837 | 40.515 |
| **M14-MUT1** | VEN+PIT | 3.105 | 17.026 | 9.651 | 28.106 |
| **M14-MUT2** | VEN+PIT | 24.591 | 37.739 | 29.146 | 46.236 |
